# Dynamical Responses Predict a Distal Site that Modulates Activity in an Antibiotic Resistance Enzyme

**DOI:** 10.1101/2024.04.29.591639

**Authors:** Michael Beer, Ana Sofia F. Oliveira, Catherine L. Tooke, Philip Hinchliffe, Angie Tsz Yan Li, Balazs Balega, James Spencer, Adrian J. Mulholland

**Affiliations:** School of Cellular and Molecular Medicine, University of Bristol, Bristol, United Kingdom, BS8 1TD; Centre for Computational Chemistry, School of Chemistry, University of Bristol, United Kingdom, BS8 1TH

**Author notes:** <u>Corresponding Authors</u>. Department of Biology and Biochemistry, 4 South, Claverton Down, University of Bath, BA2 7AY (Catherine L. Tooke). Li Ka Shing Faculty of Medicine, Hong Kong, 30-32 Ngan Shing St, Sha Tin, University of Hong Kong, Hong Kong (Angie Tsz Yan Li).

## Abstract

β-Lactamases, which hydrolyse β-lactam antibiotics, are key determinants of antibiotic resistance. Predicting the sites and effects of distal mutations in enzymes is challenging. For β-lactamases, the ability to make such predictions would contribute to understanding activity against, and development of, antibiotics and inhibitors to combat resistance. Here, using dynamical non-equilibrium molecular dynamics (D-NEMD) simulations combined with experiments, we demonstrate that intramolecular communication networks differ in three class A SulpHydryl Variant (SHV)-type β-lactamases). Differences in network architecture and correlated motions link to catalytic efficiency and β-lactam substrate spectrum. Further, the simulations identify a distal residue 89 in the clinically important *Klebsiella pneumoniae* carbapenemase 2 (KPC-2), as a participant in similar networks, suggesting that mutation at this position would modulate enzyme activity. Experimental kinetics, biophysical and structural characterisation of the naturally occurring, but previously biochemically uncharacterised, KPC-2^G89D^ mutant with several antibiotics and inhibitors reveals significant changes in hydrolytic spectrum, specifically reducing activity towards carbapenems without effecting major structural or stability changes. These results show that D-NEMD simulations can predict distal sites where mutation affects enzyme activity. This approach could have broad application in understanding enzyme evolution, and in engineering of natural and *de novo* enzymes.

## Main

Antimicrobial resistance (AMR) is a growing global healthcare crisis, associated with 4.95 million deaths in 2019 [1]. β-lactams account for approximately 65% of all antibiotic usage in humans worldwide [2] and are vital components of our antibiotic arsenal. There are four major β-lactam classes: penicillins, cephalosporins (including oxyiminocephalosporins such as ceftazidime), carbapenems, and monobactams (Figure S1). In Gram-negative bacteria (such as *E. coli*), which are leading causes of antibiotic-resistant infections worldwide, the primary mechanism of β-lactam resistance is expression of β-lactamases [2], enzymes that hydrolyse the β-lactam ring to abolish antibacterial activity [3] (Figure 1). There are four classes (A-D) of β-lactamases, with class A being the largest and most widely disseminated [3]. This includes numerous enzyme groups, including the widely distributed SHV (SulpHydryl Variant) and KPC (*Klebsiella pneumoniae* carbapenemase) families, and collectively has activity against all clinically used β-lactam antibiotics [4]. Single and multiple amino acid substitutions expand the β-lactamase spectrum of activity to cover new β-lactams and/or reduce susceptibility to β-lactamase inhibitors that are co-administered with β-lactams to treat resistant infections [5]. While, in some cases, it is clear that individual mutations exert their effects by altering active site structure [6], others are situated far from the active site and the reasons for their (often profound) effects upon activity are obscure. Understanding and predicting the effects of β-lactamase mutations will inform more effective β-lactam use and drive the development both of new β-lactam antibiotics and of more effective β-lactamase inhibitors.

**Figure 1:**
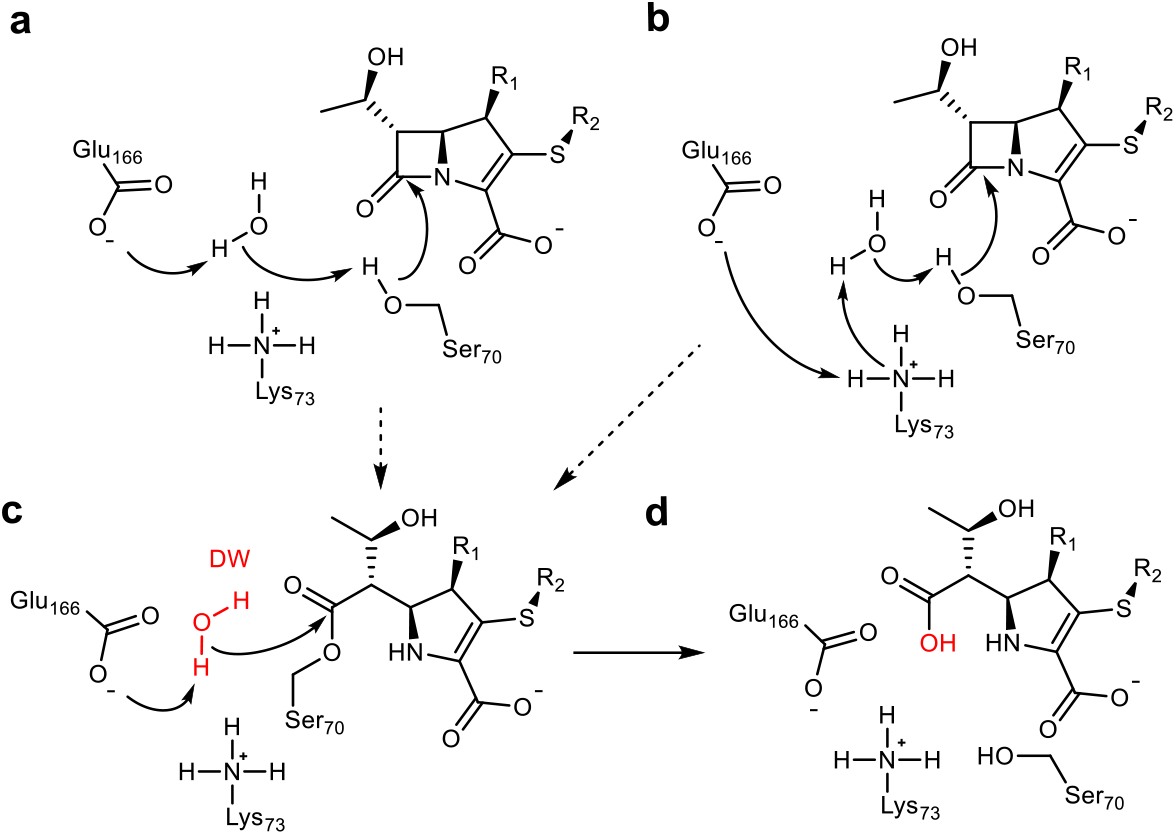
β-Lactam hydrolysis catalysed by class A β-lactamases. Substrate shown is a generalised carbapenem. Acylation (dashed arrows) catalysed by Glu166 (a) or Glu166/Lys73 (b) general bases forms the covalent acyl-enzyme (c). Deacylation requires nucleophilic attack of the deacylating water molecule (DW, red), activated by proton transfer to Glu166, on the acyl-enzyme carbonyl, to regenerate the enzyme and liberate the hydrolysed β-lactam (d) [3].

Remote mutations are well known to influence the behaviour of both natural and designed enzymes [7, 8], yet understanding of structure and function is still largely concentrated on active sites and binding interfaces. Understanding the roles of residues in distal regions will expand the scope of rational protein design [8]. Furthermore, the ability to effectively predict and identify the impact of mutations remote from active sites and ligand/protein binding regions should aid therapeutic development for multiple clinical pathologies e.g. cancer [9], HIV/AIDS [10, 11] and mental health disorders [12]. There have been previous attempts to employ path calculation methods to interrogate the conformational ensembles of proteins and also to predict residues key to catalysis, through the analysis of equilibrium simulations or static PDB structures [13-15]. Here, we employ dynamical non-equilibrium molecular dynamics (D-NEMD) simulations, an emerging computational technique that applies the Kubo-Onsager relation [16, 17] to measure the linear response of a protein to a perturbation that pushes the system out of equilibrium. These simulations can reveal allosteric communication networks within proteins [18-25]. In D-NEMD, an external perturbation (e.g. deletion of a bound ligand) is applied to an equilibrium simulation. This enables the response of the protein to the perturbation to be directly measured by comparing the equilibrium trajectory with multiple parallel non-equilibrium simulations (Figure S2) [25]. This conceptually simple but powerful approach enables computing of the time-dependent dynamic response of the protein, with assessment of statistical significance. D-NEMD simulations have recently been applied to identify structural communication pathways in a range of biomolecular systems [18, 20, 21, 24-28]. D-NEMD simulations of the β-lactamase enzymes KPC-2 and TEM-1 previously deleted an allosterically bound ligand as the applied perturbation, identifying a network of residues that link the allosteric site to the active site. Here, we apply D-NEMD to show that such networks differ between point variants of the SHV β-lactamase with diverse activities towards different β-lactam substrates; and to identify a site in the KPC-2 β-lactamase where we would predict mutation to affect activity through communication with the active site. Using steady state and pre-steady state kinetics, circular dichroism spectroscopy and high-resolution X-ray crystallography, we reveal that a single predicted mutation, situated within a distal loop, significantly impacts the KPC-2 activity spectrum. This work highlights the effectiveness of D-NEMD simulations as a method to identify previously uncharacterised mutations, distant from enzyme active sites, that affect activity.

### Communication Networks Differ Between Single-Point Variants of SHV β-Lactamases

The SHV family of class A β-lactamases includes enzymes with broad-spectrum, extended-spectrum and carbapenemase activity [29]. Broad β-lactamases efficiently hydrolyse penicillins, the most commonly prescribed antibiotics in the UK, and some early generation cephalosporins [30]. Extended spectrum β-lactamases (ESBLs) further hydrolyse the later generation oxyiminocephalosporins, widely used antibiotics that are on the World Health Organization’s list of essential medicines. Carbapenemases turnover carbapenem antibiotics, previously considered ‘last resort’ β-lactams. Many single-point variants of the broad-spectrum parent enzyme, SHV-1 (Figure 2a) have altered activity profiles. For example, SHV-2 (SHV-1^G238S^) and SHV-38 (SHV-1^A146V^) have increased activity against oxyiminocephalosporins and carbapenems, classing them as extended-spectrum and carbapenemase enzymes, respectively (Table S1) [29, 31]. We used D-NEMD simulations to investigate how single-point variations may affect internal communication networks in SHV enzymes. We compare the structural and dynamic responses of SHV-1, SHV-2 and SHV-38 to the removal of a non-covalent active site ligand (sulbactam, Figure S1). Cα deviations after 5ns of non-equilibrium simulation were compared with the equivalent time points of the unperturbed system, averaged over 200 non-equilibrium simulations (5 equilibrium simulations (Figure S2, S3) with non-equilibrium simulations started from snapshots every 5 ns from 50 ns to 250 ns). These deviations reveal communication networks that differ between the variants (Figures 2b, c, S4).

**Figure 2:**
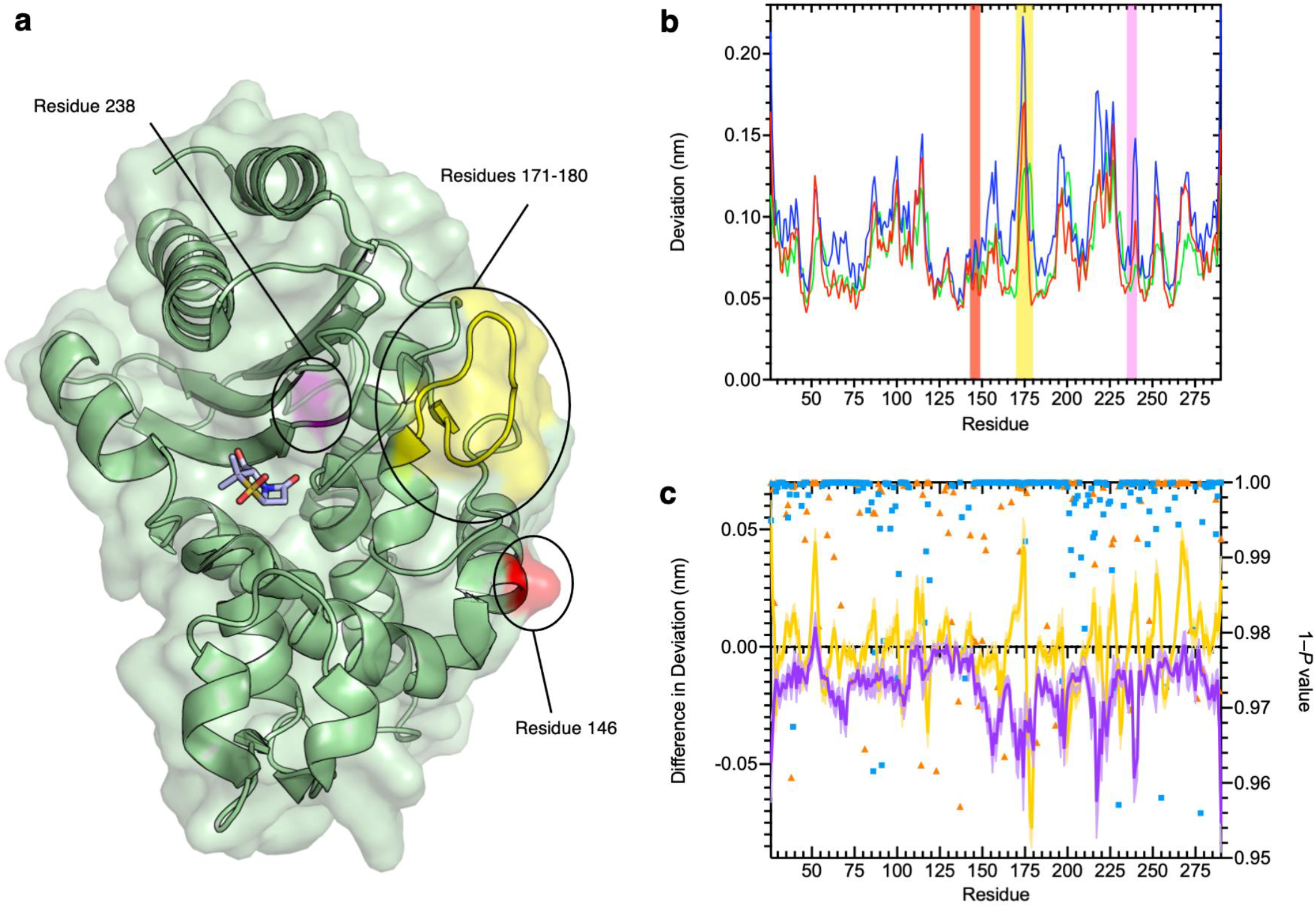
Residue deviations in SHV-1, -2 and -38 from D-NEMD simulations. a) Cartoon of SHV-1 β-lactamase, showing positions of mutations in SHV-2 (SHV-1 G238S, pink) and SHV-38 (SHV-1 A146V, red), and part of the Ω-loop (residues 171-180, yellow) close to the active site. b) Per-residue deviations calculated using the Kubo-Onsager relation [16, 17] for SHV-1 (red), SHV-2 (blue) and SHV-38 (green). Residues 146 (red bar), 171-180 (yellow bar) and 238 (pink bar) are highlighted. Deviations are calculated by averaging Cα RMSD values between perturbed (i.e. after removal of bound sulbactam ligand) and unperturbed systems 5 ns after perturbation for each residue c) Differences in deviations 5ns after perturbation, highlighting the effect of mutations on the communication network. Plots show SHV-1 vs. SHV-2 (purple) and SHV-1 vs. SHV-38 (yellow). Negative values indicate that SHV-2 or SHV-38 have greater Cα deviations than SHV-1 at specified residues. Error bars (one standard error of difference) are displayed above and below difference plots (lighter shading). Symbols show 1 – *P* values ≥ 0.95 (right axis, indicating statistical significance) for differences in deviation for SHV-1 vs. SHV-2 (blue squares) and SHV-1 vs. SHV-38 (orange triangles).

The networks identified by the D-NEMD Cα deviations show connections of multiple regions to the active site in all three SHV variants (Figure S5). For all three enzymes, the α8-α9 loop (residues 193-200) and α10 (residues 215-230) helix and the Ω-loop (171-180) all show significant D-NEMD Cα deviations, with moderate deviations also evident for the α3 and α4 helices (Figure S5). However, both SHV-2 and SHV-38 show different structural responses to the perturbation, compared to SHV-1. The average difference in deviation between SHV-1 and SHV-2 is 0.02 nm, and between SHV-1 and SHV-38 is –0.0005 nm. The response of SHV-2 is greater than that of either SHV-1 or SHV-38 in multiple regions, including the Ω-loop (residues 171-180) that borders the catalytic site (Figures 2a, b, S4, S5). SHV-2 therefore has more tightly correlated movements, and is more responsive to active site perturbation, than either SHV-1 or SHV-38. Our data indicate that the G238S substitution in SHV-2, that is considered to facilitate cephalosporin hydrolysis through local expansion of the active site, also exerts a more generalised effect upon enzyme dynamics via the intramolecular network [32]. While SHV-1 and SHV-38 have similar response amplitudes, there are multiple significant differences at specific positions. For example, there is a large deviation at residue 175 in SHV-1, whereas the corresponding peak in SHV-38 is at residue 178. Hence, while the same regions are involved in the communication networks within each SHV enzyme, the precise architecture of each network differs between variants (Figure 2c, S4, S5).

The results presented here indicate that individual point mutations influence the global dynamic behaviour of the enzyme through changes to correlated motions. In turn, these changes in dynamical networks can apparently affect activity. The results indicate that both the precise architecture of such networks (i.e. the locations of nodes showing statistically significant deviations between equilibrium and non-equilibrium simulations) reflect activity towards both specific types of β-lactam substrate (i.e. oxyiminocephalosporins for the extended-spectrum SHV-2 enzyme and carbapenems in the case of SHV-38). Such networks provide a mechanism by which remote regions of proteins are connected to regions of direct functional interest, such as the active site, with the implication that changes within them can modulate activity.

### Predicting sites of mutation that affect enzymatic activity

The KPC-2 β-lactamase is an enzyme that efficiently hydrolyzes carbapenems, the most potent β-lactam antibiotics, active against the broadest range of Gram-positive and Gram-negative bacteria of any β-lactam class [33]. Previous D-NEMD simulations revealed a network of residues that connect an allosteric ligand binding site to the KPC-2 active site region, providing first indications that these simulations can identify distal regions that modulate enzyme activity [28]. We use D-NEMD simulations here to identify regions significantly affected by a perturbation a removal of an orthosteric ligand (heteroaryl phosphonate compound 2 [34], Figure S1): such regions potentially contain sites distant from the active site that affect activity (either the turnover or substrate spectrum). To verify the calculated networks were not just capturing random equilibrium motions, a comparison of the networks calculated by D-NEMD simulations to a control simulation of a ‘null’ perturbation (randomisation of atomic velocities within the Boltzmann distribution) was performed (Figure S6).

KPC-2 and SHV β-lactamases have highly similar global structures (RMSD of 1.5 Å, over 241/265 Cα atoms). However, the networks calculated after active site ligand removal differ between the two enzymes. Deviations are observed in the KPC-2 α2-β4 (residues 81–93) and α11-β7 (residues 226–236) loops, but neither region showed prominent deviations in the SHV variants. The participation of the α2-β4 loop in the dynamical network in KPC-2 indicates strongly correlated behaviour between this distal region of the protein and the active site despite being relatively far apart (26.3 Å distance between Cα atoms of the catalytic Ser70 and Gly89 in the α2-β4 loop).

Searches of the β-lactamase database [31] identified the naturally occurring KPC variant KPC-59 as containing a single mutation within the α2-β4 loop (glycine to aspartate substitution at position 89 (G89D) compared to the parent KPC-2 enzyme); but kinetic characterizations of KPC-59 are so far unreported. Accordingly, the properties of KPC-2^G89D^ were investigated by both D-NEMD simulations and experimental kinetic and structural characterisation of the purified recombinant enzyme.

Models of KPC-2^G89D^ and its complex with compound 2 were first generated using ColabFold [35] and modelling into KPC-2 active site electron density. For D-NEMD, 200 non-equilibrium simulations of KPC-2^G89D^ were performed, removing compound 2 from the active site, and the responses were compared against results for the same perturbation in KPC-2 (Figure 3). Multiple statistically significant changes in the response amplitudes of specific regions KPC-2 and KPC-2^G89D^ are observed (Figure 3b, S8): e.g., at position 234 (a highly conserved residue involved in the catalytic mechanism of KPC-2 and other class A β-lactamases) [36-38]. The magnitude of the differences in deviations between KPC-2 and KPC-2^G89D^ are smaller than those between SHV variants (Figure 2), consistent with the greater rigidity and stability of KPC enzymes compared to other class A β-lactamases [39]. The differences in calculated residue networks indicate that a glycine to aspartate substitution at position 89 affects the intramolecular communication network within KPC-2. This provides support for the initial D-NEMD prediction (above) that the G89D mutation affects activity.

**Figure 3:**
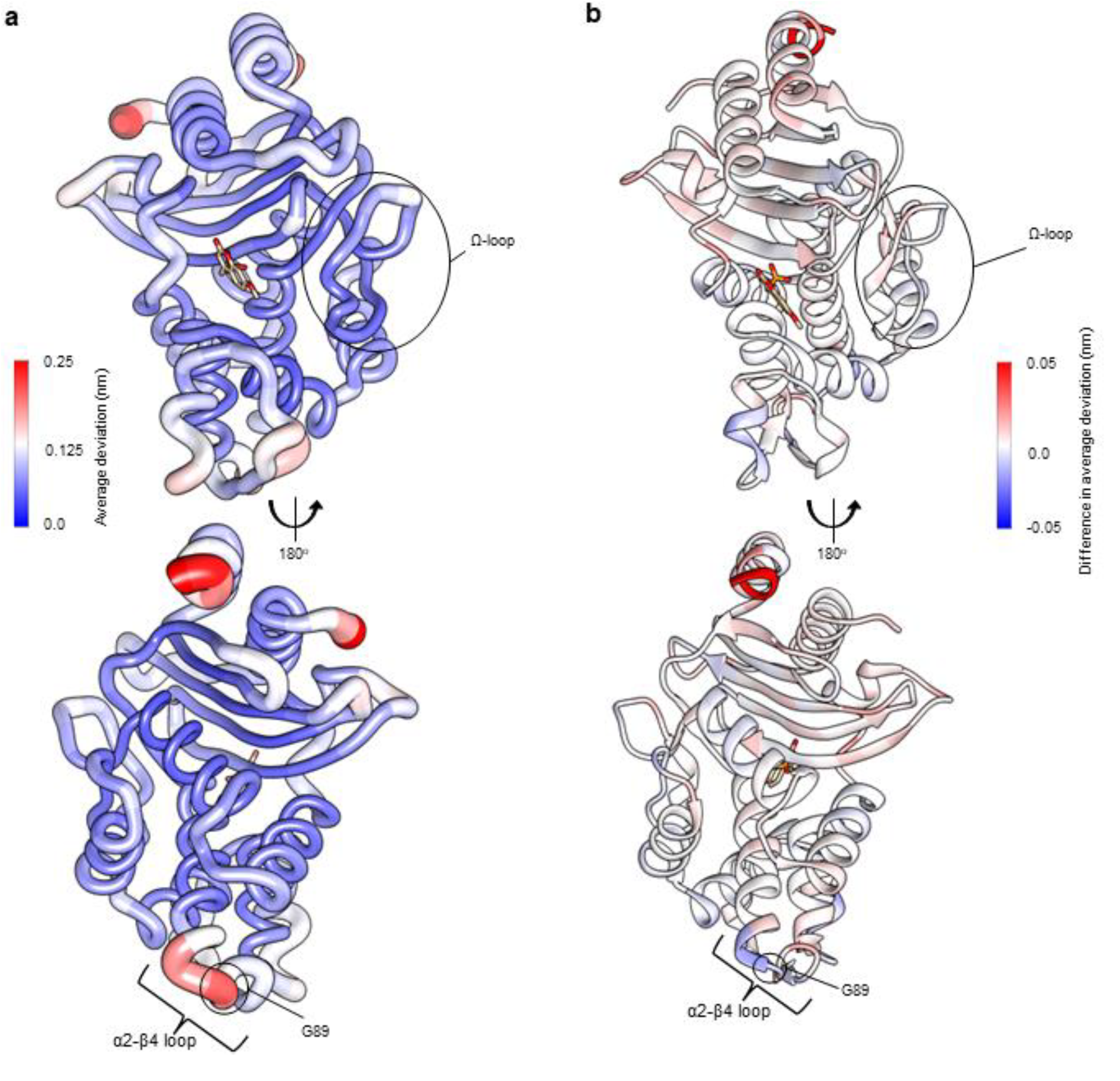
Residue deviations for KPC-2 and KPC-2^G89D^ from D-NEMD simulations. Deviations are calculated from the differences in Cα positions between perturbed and unperturbed systems (nonequilibrium vs equilibrium simulations at equivalent time points) for each residue using the Kubo-Onsager relation [16, 17]. Cα deviations were then averaged over all 200 simulations (non-equilibrium simulations started from snapshots of 5 equilibrium simulations taken every 5 ns from 50 ns to 250 ns, Figures S2 and S7). a) Residue deviations for KPC-2 5ns after deletion of the active site ligand rendered onto KPC-2 crystal structure (PDB ID 6D16 [34]). b) Difference in average residue deviations between KPC-2 and KPC-2^G89D^ 5ns after deletion of the active site ligand rendered onto KPC-2 crystal structure (PDB ID 6D16). Positive values (red) indicate where Cα deviations were greater in KPC-2, negative values (blue) indicate where Cα deviations were greater in KPC-2^G89D^. Many difference values are statistically significant, due to the large number of replicates obtained using the D-NEMD approach (Figures S2 and S8). The KPC^G89D^ structure was created using the ColabFold tool [35] (Figure S9). The active site ligand (compound) is shown in stick form to highlight the active site region.

To test this prediction, the KPC-2^G89D^ mutant was generated and purified from recombinant *E. coli*. Circular dichroism (CD) spectroscopy thermal melting experiments on the purified mutant protein indicated that the mutation does not adversely affect global stability, conferring a slight (1.1 °C) increase in melting temperature (*T*^m^) (Figure S10).

Strikingly, steady-state kinetic measurements (Table 1) reveal significant changes to the hydrolytic profile of KPC-2^G89D^, compared to that of wild-type KPC-2. Notably, while KPC-2 has strong carbapenemase activity, the G89D mutation results in a 100-fold decrease in catalytic efficiency (*k*_cat_/*K*_M_, Table 1) towards meropenem and a 10-fold decrease for imipenem (Figure S1). There is also a 10-fold decrease in the rate of turnover of the 1^st^ generation cephalosporin cephalothin (Figure S1). This substantial decrease in carbapenem and cephalothin-hydrolysing activity is combined with an increase in catalytic rate (*k*_*cat*_, Table 1) for hydrolysis of the 3^rd^ generation oxyiminocephalosporin cefotaxime (Figure S1). Moreover, *K*_M_ values for the oxyiminocephalosporins ceftazidime and cefotaxime showed significant increases compared to the parent enzyme KPC-2. In contrast, KPC-2^G89D^ is somewhat more active against penicillin. These data show the G89D mutation to affect KPC-catalysed hydrolysis of specific β-lactam substrates, rather than exerting a general effect upon enzymatic activity. This highlights the ability of the D-NEMD approach to identify residues that modulate (and can increase some) activity, rather than those that are directly catalytic, purely structurally significant or affect global stability, for which mutation would be expected to abolish activity towards all substrates.

**Table 1:** Steady-state kinetic parameters for hydrolysis of range of β-lactam substrates (Figure S1) by KPC-2 and its G89D variant. Standard errors for *k*_cat_ and *K*_M_ values are shown in parentheses.

| β-Lactam Antibiotic | β-Lactam Class | $k_{cat}$ ($s^{-1}$) | | $K_M$ ($\mu M$) | | $k_{cat}/K_M$ ($s^{-1} \mu M^{-1}$) | |
| --- | --- | --- | --- | --- | --- | --- | --- |
|  |  | KPC-2<sup>G89D</sup> | KPC-2 | KPC-2<sup>G89D</sup> | KPC-2 | KPC-2<sup>G89D</sup> | KPC-2 |
| Ampicillin | Penicillin | 160<br>(5.83) | 82.3<br>(3.92) | 201<br>(36.3) | 271<br>(44.7) | 0.79 | 0.3 |
| Cephalothin | Cephalosporin | 10.8<br>(0.32) | 112<br>(2.78) | 45.1<br>(3.71) | 41.6<br>(3.41) | 0.24 | 2.68 |
| Ceftazidime | Cephalosporin | 2.7<br>(0.36) | 1.9<br>(0.12) | 1340<br>(220) | 533<br>(69) | $2 \times 10^{-3}$ | $3.5 \times 10^{-3}$ |
| Cefotaxime | Cephalosporin | 523<br>(176) | 75.8<br>(6.61) | 2470<br>(942) | 199<br>(29) | 0.21<br>(0.14<sup>a</sup>) | 0.38 |
| Meropenem | Carbapenem | 0.2<br>($1 \times 10^{-2}$) | 21.5<br>(0.12) | 7.9<br>(1.57) | 7.1<br>(1.01) | 0.03 | 3.00 |
| Imipenem | Carbapenem | 2.8 | 22 | 68.9 | 72 | 0.04 | 0.31 |

<sup>a</sup> $k_{cat}/K_M$ value calculated from analysis of complete progress curves. This was done for cefotaxime due to the large errors resulting from a high $K_M$ value.

To confirm the direct impact of the G89D mutation on β-lactam turnover, rather than binding, the interactions of KPC-2 and KPC-2^G89D^ with the carbapenem meropenem were investigated in pre-steady state kinetic assays under pseudo-first order conditions, monitoring tryptophan fluorescence as previously performed with OXA-48 β-lactamase [40]. These experiments showed that the forward and reverse rate constants *k*_1_ and *k*_-1_ for meropenem binding, and the derived *K*_D_ values, (3.7 and 5.9 μM for KPC-2 and KPC-2^G89D^, Figure S11), are similar for the two enzymes, indicating that the G89D substitution does not affect meropenem binding. This shows that the 100-fold decrease in activity caused by this mutation is likely due to a change in the rate of the reaction on the enzyme, suggesting the G89D substitution has an impact on the active site chemistry and thus is a catalysis modulating mutation.

Point variants of β-lactamases can also affect the efficacy of mechanism-based inhibitors that are used clinically in combination with susceptible β-lactams [41] either through changes in binding interactions or changes in turnover capability [5, 42, 43]. The combination of ceftazidime with the reversible, diazabicyclooctane (DBO) inhibitor avibactam is effective against most KPC-producing organisms, which generally evade the action of mechanism-based β-lactam inhibitors such as clavulanic acid. However, KPC variants are now emerging that reduce susceptibility of producer organisms to ceftazidime-avibactam. Accordingly, we investigated the *in vitro* potency of selected DBO inhibitors against KPC-2^G89D^. No significant change was observed in the IC_50_ value of avibactam (Figure S1) against the KPC-2^G89D^ mutant compared to the wild-type enzyme (12.2 nM vs 10 nM for KPC-2) [44]. However, the G89D mutation decreases potency of inhibition by the bulkier DBO inhibitor zidebactam (Figure S1), with a 10-fold increase in IC_50_ value (0.7 nM vs 0.06 nM for KPC-2).

To investigate the basis of these changes in the activity spectrum of KPC-2^G89D^, we determined X-ray crystal structures of the uncomplexed enzyme (Figure S12, S13) and of its covalent complexes with the carbapenems imipenem and meropenem and with avibactam (Table S2). The crystal structure of uncomplexed KPC-2^G89D^ revealed that, consistent with CD data, the mutation does not affect the global structure of the enzyme, (Cα RMSD 0.07 Å to uncomplexed KPC-2, PDB ID 5UL8 [45]) (Figure 4a, b, SI Note 1).

**Figure 4:**
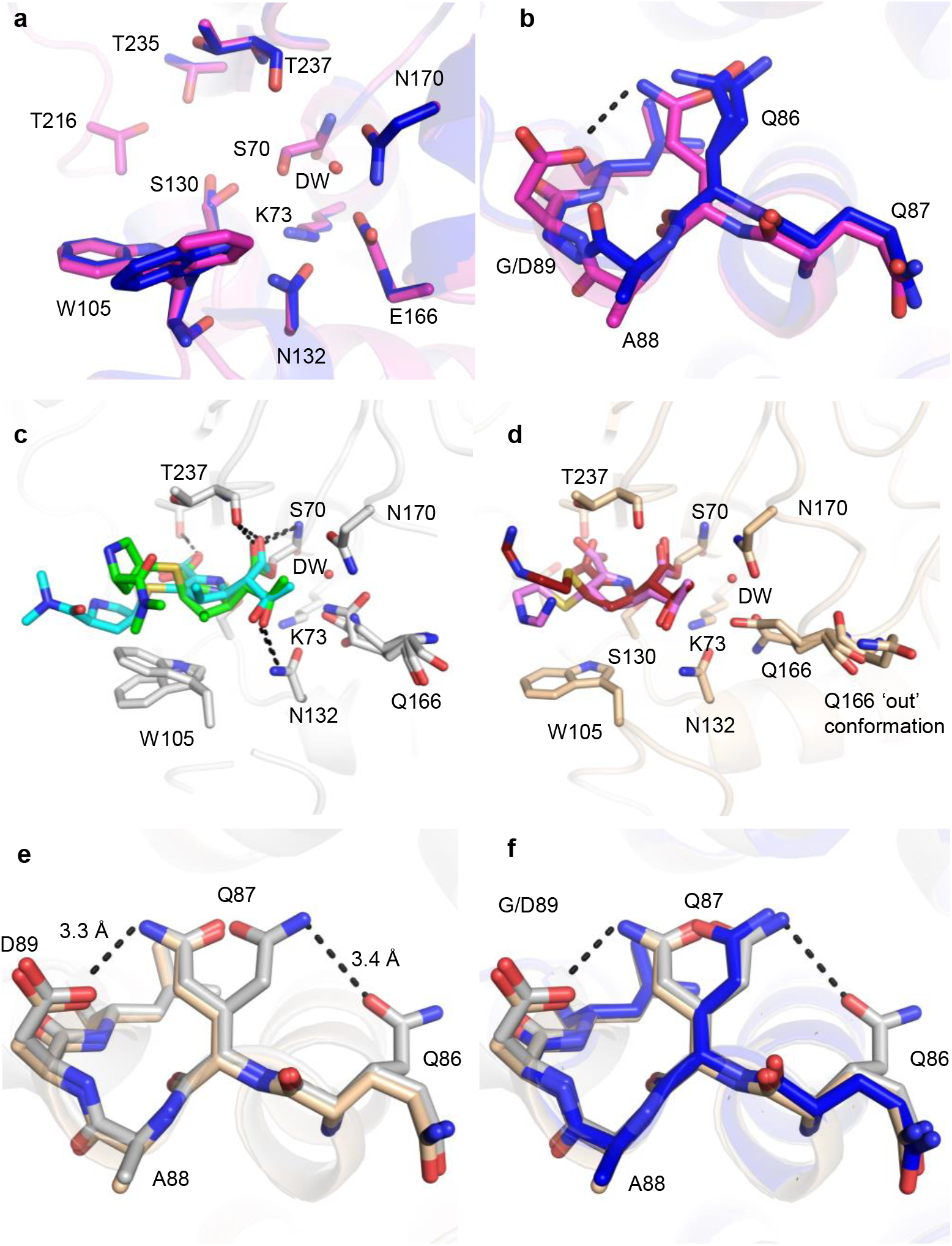
Crystal structures of the KPC-2^G89D^ mutant compared with KPC-2. a) Active sites of uncomplexed (apo) KPC-2^G89D^ (magenta, this work) and KPC-2 (PDB ID 5UL8 [45], blue). Residues implicated in catalysis are shown as sticks, and the deacylating water molecule (DW) as a red sphere. b) α2-β4 loops of KPC-2^G89D^ (magenta) and KPC-2. The additional hydrogen bond between D89 and Q87 in the KPC-2^G89D^ structure is highlighted (black, dashed line). c) Active site of the KPC-2^G89D/E166Q^:meropenem acyl-enzyme (side chain carbon atoms in grey, Δ2 tautomer in cyan and Δ1-(2*R*) in green, Figure S15, S16, S17). Hydrogen bonds between the enzyme and meropenem are shown (black, dashed lines). d) Active site of the KPC-2^G89D/E166Q^:imipenem acyl-enzyme (side chain carbon atoms tan, Δ1- (2*R*) tautomer in red and Δ1-(2*S*) tautomer in pink, Figure S14, S15, S16, S17). Hydrogen bonds are those observed in the KPC-2^G89D/E166Q^:meropenem complex (distances between atoms differ, Figure S16). Note multiple conformations of Gln166 in both KPC-2^G89D/E166Q^:carbapenem complexes, in particular the ‘out’ conformation of Gln166 in the KPC-2^G89D/E166Q^:imipenem acyl-enzyme. e) architecture of the α2-β4 loop in KPC-2^G89D/E166Q^:meropenem acyl-enzyme complex (grey) and the KPC-2^G89D/E166Q^:imipenem acyl-enzyme complex (tan). f) α2-β4 loop in the KPC-2^G89D/E166Q^:meropenem derived acyl-enzyme complex (grey) and the KPC-2^G89D/E166Q^:imipenem derived acyl-enzyme complex (tan) and KPC-2:meropenem (PDB 8AKL, blue) [46].

To capture the structures of carbapenem-derived acyl-enzyme complexes, and thus probe why the G89D substitution reduces catalytic efficiency for carbapenems, the isosteric Glu166 to Gln166 substitution was made in KPC-2^G89D^ and the purified recombinant protein crystallised and challenged with imipenem and meropenem. The E166Q substitution renders class A β-lactamases deacylation-deficient but does not change their global structure (Figure S13) or prevent acylation by β-lactams [47]) [46, 48]. Diffraction data resolved meropenem- and imipenem-derived acyl-enzyme complexes with KPC-2^G89D/E166Q^ to high resolution (1.09 Å and 1.03 Å, respectively, Table S2, Figure S16, S17). Carbapenem acyl-enzymes are known to tautomerise through migration of the double bond in the 5-membered pyrroline ring, making possible observation of multiple tautomers and stereomers in the same crystal structure (SI Note 2, Figure S16, S17) [49, 50]. Accordingly, the KPC-2^G89D/E166Q^ meropenem acyl-enzyme was modelled in the Δ1-(2*R*) and Δ2 configurations, while the imipenem-derived complex was modelled as the Δ1-(2*R*) and Δ1-(2*S*) forms. We have previously shown that the different tautomers have different propensities to deacylate, with the initial state (Δ2) being more deacylation competent than Δ1 stereomers [49]. There is no difference in the orientations of either imipenem or meropenem derived acyl-enzyme KPC-2^G89D/E166Q^ complexes compared to the previously deposited meropenem and imipenem derived KPC-2^E166Q^ acyl-enzyme complexes (PDB ID 8AKL, 8AKK [49], Figure S18). Overall, analysis of the KPC-2^G89D/E166Q^ carbapenem derived acyl-enzyme complexes indicates that neither the active site architecture, nor covalent binding of carbapenem substrates, is not affected by the G89D mutation.

In both KPC-2^G89D/E166Q^:carbapenem derived acyl-enzyme structures reported here, residues (165-170) in the catalytically essential Ω-loop are highly flexible, shown e.g. by multiple conformations of Gln166 and high B-factors (Figure S19, S20). The multiple conformations of Gln166 include an ‘out’ conformation (Figure 4d, S20) in the imipenem-derived acyl-enzyme complex, where the Gln166 side chain faces bulk solvent rather than the bound carbapenem. This ‘out’ conformation of residue 166 is adopted when KPC-2 forms acyl-enzyme complexes with substrates that deacylate poorly [48, 51]. Multiple studies of a range of class A β-lactamases show that the conformational stability of the Ω-loop in β-lactam derived acyl-enzyme complexes correlates strongly with the ability to turn over that substrate [48, 49, 52, 53], suggesting that the high mobility of the Ω-loop contributes to the reduced activity of KPC-2^G89D^ towards carbapenems. B-factor (adjusted, B’-factor) analysis of KPC-2 acyl-enzyme complexes with different β-lactam substrates from previous studies highlight a negative correlation between the stability of the α2-β4 loop (containing residue 89, Figure 3) and the stability of the Ω-loop (Figure S21), i.e. structures with high B’-factors for the α2-β4 loop have low B’-factors for the Ω-loop [45, 48, 49]. In both acyl-enzyme structures of KPC-2^G89D^ presented here, the α2-β4 loop forms extra hydrogen bonds, between residues D89 and Q87, that are not observed in equivalent KPC-2 (Figure 4e, f). These hydrogen bonds result in lower B’-factors for residues in the α2-β4 loop, which combined with the observed instability/high mobility of the Ω-loop, is consistent with the observed inverse correlation between the two loops (SI Note 2, Figure S21).

A water molecule is positioned for deacylation in the resolved both crystal structures of carbapenem derived acyl-enzyme complexes with KPC-2^G89D/E166Q^ (DW, Figure 1, 4c-d). A deacylating water molecule is required to deacylate the covalent complex formed between antibiotics and class A β-lactamases [6, 49, 54, 55]. Appropriate positioning and orientation of the DW, e.g. through interaction with the side chains of residues 166 and 170 (Figure 4, S16) promotes turnover of β-lactam acyl-enzymes [56, 57]. The high resolution of the acyl-enzyme complexes here allows for refinement of occupancies. While the meropenem-derived KPC-2^G89D/E166Q^ acyl-enzyme complex contains a water molecule in the DW position, it is refined at low occupancy (0.58) compared to the structure of the parent KPC-2^E166Q^:meropenem at similar resolution (occupancy 1.00, PDB 8AKL [46]). In contrast, in the imipenem-derived KPC-2^G89D/E166Q^ acyl-enzyme complex (a substrate that KPC-2^G89D^ has an increased catalytic rate against (Table 1)), the DW could be refined at full occupancy (1.00) with a lower B-factor (Table S3). Thus, a further consequence of the increased flexibility of the Ω-loop in acyl-enzymes of KPC-2^G89D^ may be a reduction in the stability/occupancy of the deacylating water molecule, and consequent impaired deacylation. However, this effect appears to be substrate-dependent and does not appear to contribute to the observed poor imipenemase activity (Table 1), as the KPC-2^G89D/E166Q^ imipenem derived acyl-enzyme complex has full occupancy and a lower DW B-factor (19.74 vs 24.49 in the KPC-2^G89D/E166Q^ meropenem derived acyl-enzyme complex).

We also determined the crystal structure of the covalent complex of KPC-2^G89D^ with the DBO inhibitor avibactam. Consistent with previous studies, bound avibactam is observed as a mixture of the sulphated and desulphated forms (Figure S22, S23, S24) [58] [59]. Structural comparisons showed the orientations of bound avibactam, and its interactions with the KPC-2^G89D^ active site, to be near identical to the wild-type structure, consistent with the minimal difference in IC_50_ values (Figure S23). These data highlight the selective impact of the G89D substitution on activity of KPC-2 with substrates/inhibitors, further strengthening the conclusion that networks identified by D-NEMD contribute to specificity as well as overall catalytic activity.

## Conclusions

Here, we combine simulations and experiments to identify functionally important differences in the dynamics of β--lactamase enzymes. We identify distal mutation sites that affect activity and validate this prediction by experiment. D-NEMD simulations identify communication networks that differ between SHV variants. The extended-spectrum SHV-2 variant, with efficient activity against the broadest range of substrates, shows the greatest pre-residue deviations. The precise architecture of intramolecular networks may reflect differences in the activity spectrum of the enzyme variants against specific substrates, as evidenced by differences in per-residue deviations between the SHV-1 and SHV-38 variants, which differ in activity towards carbapenems. Second, the communication networks identified by D-NEMD identify sites in β-lactamase enzymes that affect activity, in particular residues remote from the enzyme active site. Simulations of KPC-2 identified residue 89 as a participant in an allosteric network that connects distal regions of the protein to the active site [22, 28]. Experimental characterisation of the (previously unstudied) KPC2^G89D^ variant substantiates this conclusion and supports the importance of intramolecular [22, 28]. communication networks for activity: we show that this mutation selectively changes activity towards specific β-lactams, in particular reducing carbapenem hydrolysis.

The data presented here demonstrate that the D-NEMD technique can identify distal positions that affect turnover of specific β-lactams. Such information may help to predict possible regions where point mutations on KPC-2 and other class A β-lactamases may occur. This significantly extends previous evidence that identified networks linking a putative allosterically binding ligand to the TEM-1 and KPC-2 active sites, indicating that these networks module catalytic activity of the respective enzymes. This knowledge should also guide antibiotic and inhibitor drug development aimed at pre-empting and overcoming resistance, potentially by targeting regions that form significant nodes in intramolecular communication networks with allosteric inhibitors. The D-NEMD approach should be a valuable tool to identify dynamic intramolecular interactions that affect protein structure and function, likely to find application in areas including drug discovery, rational enzyme design and *de novo* protein design.

## Author Contributions

MB, ASFO, JS and AJM conceived the experiments. MB performed all simulations and wet lab experiments. ASFO provided supervision for the simulations and aided in data interpretation. CLT provided supervision in site directed mutagenesis, protein purification and crystal trials and contributed to data interpretation. PH contributed to data interpretation and supervision of wet lab experiments. YTAL assisted in site-directed mutagenesis, protein purification, crystal trials, and steady-state kinetics experiments as part of an undergraduate research project. MB wrote the manuscript with input and edits from ASFO, CLT, PH, JS and AJM.

## Acknowledgments

The authors thank Marc W. van der Kamp for discussion about the work and providing feedback. All simulations were conducted using the facilities of the Advanced Computing Research Centre at the University of Bristol (http://www.bris.ac.uk/acrc/). The authors thank the Diamond Light Source (beamline I03, proposals 23269 and 31440) and the beamline scientists for their service that enabled the X-ray diffraction data presented here to be collected.

## Data Availability

Input files and ligand parameters for all conducted simulations will be made available at the University of Bristol Research Data Repository (https://data.bris.ac.uk/).

