## Supplementary Information for "Dynamical Responses Predict a Distal Site that Modulates Activity in an Antibiotic Resistance Enzyme"

[*]Corresponding Authors

### Model Set Up

For SHV-1, the crystal structure of the S70C mutant bound to sulbactam (PDB ID 4FH2[1]) was used with the Cys70 residue mutated back to Ser70 in WinCoot 0.9.8.1 and modelling into Serine 70 density from the apo structure of SHV-1 (PDB ID 1SHV[2]). The SHV-2 and SHV-38 structures were generated using AlphaFold Colab[3] and the sulbactam structure modelled into the active site region, using the SHV-1 S70C electron density. Protonation states were assigned using the PropKa program at pH 7.4[4].

The KPC-2$^{G89D}$ mutant structure was created using AlphaFold Colab[3], and the compound 2 ligand modelled into the active site using the electron density map of the compound 2:KPC-2 complex (PDB ID 6D16[5]) in WinCoot 0.9.8.1[6].

All complexes were set up for molecular dynamics simulations using GROMACS 2019.1[7] using the Amber ff14SB forcefield[8] for the protein and the GAFF parameters for the ligands[9]. Ligand atoms were parameterised using the ACPYPE server[10] and RESP charges calculated using the R.E.D. server[11]. Each system was solvated using TIP3P[12] water in a cubic box with 10 Å distance between the edge and the solute. $Na^+$ and $Cl^-$ counter ions to a concentration of 180 mM were added to neutralise the system. Each system was minimised by steepest-descent for 10000 steps, to relieve bad contacts. The system was initially equilibrated in the NVT ensemble over 500 ps with restraints on all heavy atoms (force constant of 1000 kJ $mol^{-1}$ $nm^{-1}$), using V-rescale temperature coupling. Two coupling groups were used (protein and ligand, water and ions) with a coupling constant of 0.05 ps. The next step of equilibration was performed for 500 ps in the NPT ensemble using Berendsen pressure coupling, V-rescale temperature coupling (same two coupling groups as NVT equilibration but with a coupling constant of 0.1 ps). All Cα atoms and ligand heavy atoms were restrained in this equilibration stage with a force constant of 1000 kJ $mol^{-1}$ $nm^{-1}$. A second 500 ps NPT ensemble equilibration was then performed using Berendsen pressure coupling[13] with restraints only on ligand heavy atoms (100 kJ $mol^{-1}$ $nm^{-1}$ force constant) and the same V-rescale temperature coupling parameters as the previous NPT equilibration step. A 250 ns production simulations were run in the NPT ensemble using Parrinello-Rahman pressure coupling[14] and the same V-rescale temperature coupling parameters. In the production simulations, the neighbours list was updated every 40 steps. All hydrogen bonds were constrained to their equilibrium lengths with the LINCS algorithm except for the water molecules, which were kept rigid with the SETTLE algorithm.

Five 250 ns equilibrium MD simulations each were run in the GROMACS 2019.1 software package with the prepared SHV-1, SHV-2, SHV-38, KPC-2 and KPC-2$^{G89D}$ systems[7]. The simulations were considered fully equilibrated for the D-NEMD approach after 50 ns (Figures S1-S2)

Dynamical nonequilibrium MD (D-NEMD) Simulations

To study signal propagation between the active site and the rest of the protein, 200, 5 ns long, dynamical-nonequilibrium simulations were performed for SHV-1, SHV-2, SHV-38, KPC-2 and KPC-2$^{G89D}$. These simulations drive, and allow for the characterisation of, rapid conformational changes in the system and permit mapping of the communication networks within the proteins using the Kubo-Onsager approach[15]. At each 5 ns time point, from 50 ns to 250 ns, a structure file of the entire system was extracted, resulting in 40 conformations per replicate and thus 200 starting conformations per system. The ligand atoms (bound to the active site of each system) were then deleted. The resulting non-equilibrium system was run for 5 ns in the GROMACS 2019.1 software package in identical conditions to the 250 ns MD simulations described above.

The distance between the two systems at equivalent time points (e.g. 55 ns on the equilibrium MD simulation, compared to the last frame of the 5 ns non-equilibrium MD trajectory obtained starting from the conformation extracted at the 50 ns time point on the equilibrium MD simulation) was calculated for each Cα in the protein. This was done for each system after 10 ps, 50 ps, 100 ps, 500 ps, 1 ns, 3 ns and 5 ns of simulation. Cα deviation values were averaged over all 200 non-equilibrium simulations per system and the standard deviations and standard error of the mean calculated. The Cα deviation plot for the 5 ns time points was used to identify communication networks in all systems. Significant differences in Cα deviation of individual residues between systems were calculated using the student t-test, with the cut-off set at $p$=0.05.

Protein Expression and Purification

G89D and E166Q mutations of KPC-2 were produced by site directed mutagenesis of the previously constructed pET28a-KPC-2 vector[16] The primers used to generate the E166Q mutant were 5′-TCA GCT CCA GCT GCC AGC GGT CCA G-3′ and 5′-CTG GAC CGC TGG CAG CTG GAG CTG A-3′ and site directed mutagenesis was performed using the the QuikChange II XL lightening SiteDirected Mutagenesis Kit, following the manufacurer's instructions (Agilent Genomics). Proteins were subsequently expressed and purified as previously described[17].

Enzyme Kinetics

Antibiotic hydrolysis was measured at 25°C in kinetics buffer (10 mM HEPES, pH 7.5 and 150 mM NaCl, 100 µg/mL bovine serine albumin (BSA)). Steady-state kinetic parameters were calculated by measuring the hydrolysis of β-lactam antibiotics (ampicillin $\Delta\varepsilon_{235} = -900$, cefotaxime $\Delta\varepsilon_{262} = -7660$, ceftazidime $\Delta\varepsilon_{265} = -7445$, meropenem $\Delta\varepsilon_{297} = -11{,}500$)[16-19]. Hydrolysis was followed using Greiner half area 96-well plates and a BMG CLARIOstar Plus microplate reader. GraphPad Prism 9.3.1 (GraphPad Software, La Jolla, CA, USA; www.graphpad.com) was used to calculate kinetic parameters. Initial rates of antibiotic hydrolysis measured across a range of antibiotic concentrations were used to calculate steady-state parameters according to the Michaelis-Menten equation. The *kcat*/KM value for cefotaxime was additionally calculated by fitting the complete hydrolysis curve using the following equation:

$$A_t = A_\infty + (A_0 - A_\infty)e^{-kt}$$

$IC_{50}$ values were calculated by following the initial rate of nitrocefin hydrolysis (200 µM) at 486 nM ($\Delta\varepsilon_{486}$ = 20500 $M^{-1}$ $cm^{-1}$)[16], after a 10-minute pre-incubation of enzyme and inhibitor. Both inhibitors were dissolved in kinetics buffer.

### Pre-Steady-State Kinetics

Pre-steady state kinetics were measured by mixing meropenem with KPC-2 or KPC-$2^{G89D}$ in an Applied Photophysics SX20 Stopped-Flow spectrometer connected to a photodiode array detector in kinetics buffer. Data were fitted from 0.005 to 0.3 s for both KPC-2:meropenem and KPC-$2^{G89D}$:meropenem. $k_1$ and $k_{-1}$ were calculated from the gradient and the y-intercept of the $k_{obs}$ vs substrate concentration curves. $K_D$ is calculated using the following equation:

$$K_D = \frac{k_{-1}}{k_1}$$

### Circular Dichroism (CD)

Protein samples were prepared to a concentration of 20 µM in potassium phosphate buffer (100 mM, pH 7.6). The CD spectra were obtained using a JASCO J-1500 spectrophotometer. For thermal melt experiments, CD signal was measured at 220 nm between 5 °C – 95 °C, changing the temperature 1 °C per minute. A full CD spectra at 25°C for each protein was also recorded, measuring between 200-250 nm. Eight spectra were obtained for each protein and the results averaged to give the final spectra (Figure S6). Here, we report CD results as mean residual ellipticity (MRE) which was obtained from the raw data using the equation reported in Hutchins *et al*[20].

### Crystallisation and Ligand Soaking

Crystals were grown at 20°C using sitting drop vapor diffusion in CrysChem 24-well plates (Hampton Research), with wells equilibrated against 500 uL crystal buffer (5% (v/v) ethanol with 1.8-2.0 M $NH_4(SO_4)_2$). Drops consisted of 1 uL of crystal seed (generated from crushed KPC-2 crystals), 2 µL protein (30 mg/ml aliquots), and 1 µL of crystallisation reagent[17].

KPC-2$^{G89D}$ crystals were soaked in mother liquor supplemented with 30 mM avibactam for 4 hours, before being brief exposure to mother liquor supplemented with 25% (v/v) glycerol and flash frozen in liquid nitrogen. For the meropenem acylenzyme structure, crystals were soaked in mother liquor supplemented with 60 mM meropenem for 24 hours, before being briefly soaked in mother liquor with 25% (v/v) glycerol and flash frozen in liquid nitrogen.

X-ray Diffraction Data Collection and Structural Determination

Diffraction data were collected at Diamond Light Source on beamline i03 (Table S1), using an Eiger2 XE 16M detector with an exposure time of 0.004 s per image.

In all cases the images were indexed and integrated using the Dials[21] and Xia2[22] processing pipelines at Diamond Light Source. Phases were calculated using Fourier transform in PHENIX [23] with KPC-2$^{E166Q}$ with the meropenem ligand removed (PDB ID 8AKL[24]) as the starting structure. The structure was completed with iterative rounds of refinement in PHENIX [23] and manual model building in Wincoot[6]. All ligand restraints were calculated using the Grade web server (http://grade.globalphasing.org/).

SI Note 1

#### Crystallographic Analysis of Uncomplexed KPC-2$^{G89D}$

There are no large position or orientation changes of active site residues, or in the location of the deacylating water molecule (DW, Figure 1, 4a, S12), important to β-lactam hydrolysis. However, there are small differences in the α2-β4 loop, where residue 89 is located, with the carbonyl oxygen of A88 flipped 180° (but maintaining a cis conformation) to enable an additional hydrogen bonding interaction between the side chain amide of Q87 and a side chain carboxylate oxygen of D89 in KPC-2$^{G89D}$ (Figure 4B, S12).

SI Note 2

#### Crystallographic Analysis of the KPC-2$^{G89D/E166Q}$:Carbapenem Complexes

Aside from the Ω-loop instability, the active site of both KPC-2$^{G89D/E166Q}$:carbapenem complexes closely resembles that of KPC-2$^{E166Q}$:carbapenem complexes[42]. The oxyanion hole, formed from the backbone amides of Ser70 and Thr237 is maintained, with hydrogen bonds between each amide and the carbonyl oxygen (Figures 4c, 4d, S15) present in both KPC-2$^{G89D/E166Q}$:carbapenem structures. Furthermore, a hydrogen bond interaction between the 6a-hydroxyethyl group of both carbapenems and the side chain of Asn132 is retained. This orients the hydroxyethyl group away from the deacylating water, a factor that known to promote carbapenemase activity of class A β-lactamases[9]. Additionally, in both KPC-2$^{G89D/E166Q}$:carbapenem structures, Ser130 is oriented within hydrogen bonding distance of the β-lactam amide (Figures 4c, 4d, S15), as is also seen in the KPC-$2^{E166Q}$:carbapenem complexes.

It is noted that the B'-factor of the Ω-loop in the KPC-$2^{G89D/E166Q}$:carbapenem complexes is lower than other complexes modelled with poor substrates. This is due to the multiple modelled conformations of Gln166, which prevents accurate B'-factor comparison with the KPC-$2^{E166Q}$:β-lactam structures. The multiple modelled conformations in these structures does, though, provide another means of identifying that the Ω-loop is flexible in the KPC-$2^{G89D/E166Q}$:carbapenem complexes, whilst the residue loop region is more stable than in the KPC-$2^{E166Q}$:carbapenem acyl-enzyme complex structures due to its additional electrostatic interactions.

**1**

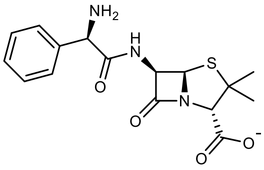

Ampicillin

**2**

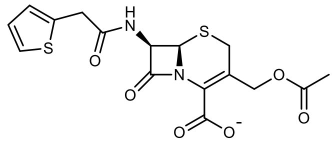

Cephalothin

**3**

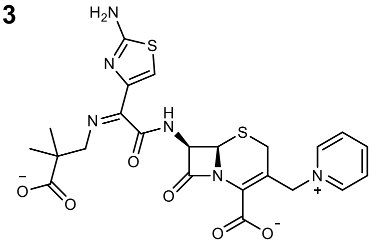

Ceftazidime

**4**

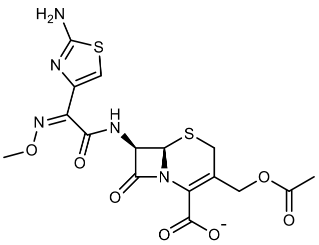

Cefotaxime

**5**

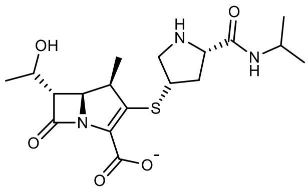

Meropenem

**6**

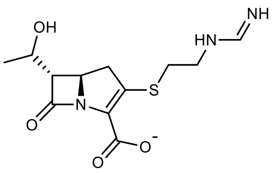

Imipenem

**7**

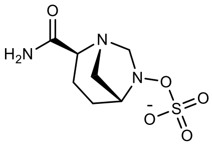

Avibactam

**8**

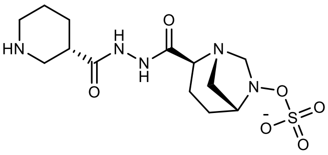

Zidebactam

**9**

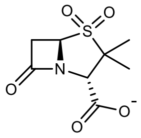

Sulbactam

**10**

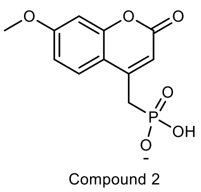

Compound 2

**Figure S1: Structures of β-lactam antibiotics and β-lactamase inhibitors.**

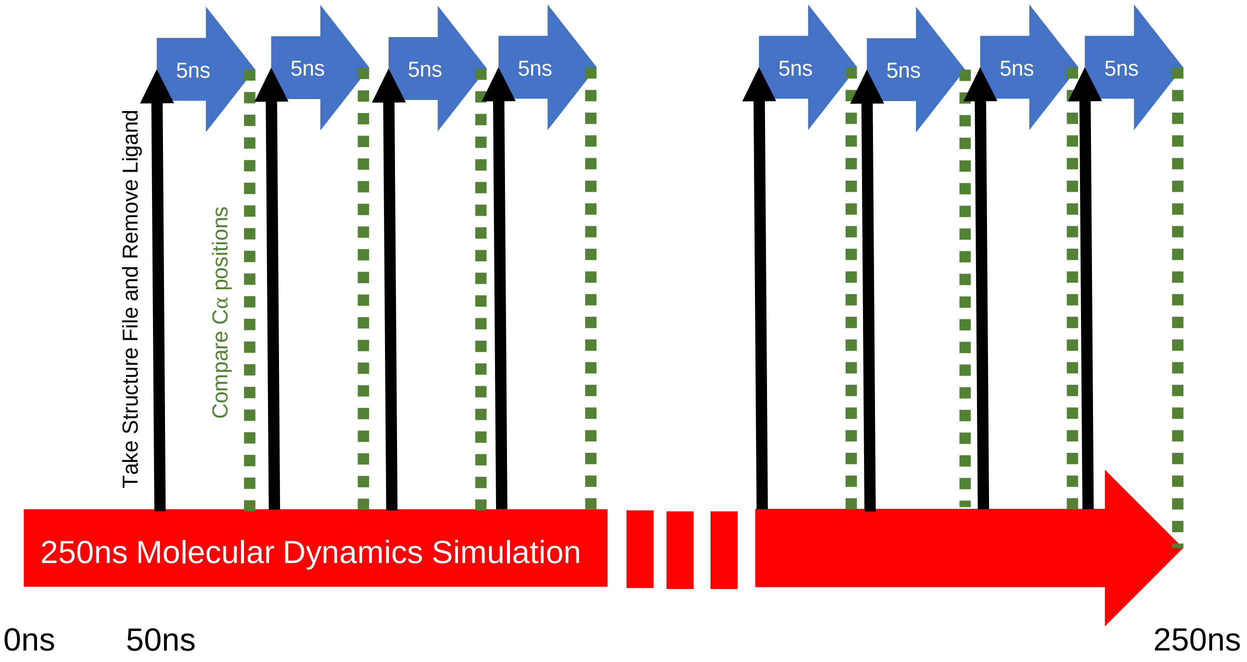


**Figure S2: Theoretical framework of dynamical non-equilibrium molecular dynamics (D-NEMD) simulations.** Large red arrow represents a 250 ns equilibrium molecular dynamics (MD) simulation e.g. of SHV-1 with sulbactam bound non-covalently to the active site. Black lines represent individual structure files written at each 5 ns time point on the equilibrium MD simulation from which the active site ligand atoms i.e. of sulbactam are removed. Blue arrows represent new non-equilibrium simulations starting from each of these structure files. Green dashed lines show comparisons of Cα positions between the non-equilibrium and equilibrium simulations at equivalent time points from which residue deviations are calculated [15]. This procedure is repeated for every 5ns of the 250 ns equilibrium simulation, from 50 ns onwards, i.e. totalling 40 repeats per equilibrium simulation. With 5 repeats for each system (1.25 µs of equilibrium simulation time) this results in a total 200 x 5 ns non-equilibrium simulations per system. A large number of repeats is often required to obtain statistically significant Cα deviations, in order to negate the impact of natural residue fluctuations during an MD simulation.

**Table S1: Kinetic parameters for SHV-1, SHV-2 and SHV-38.**

| β-Lactamase | β-Lactam Antibiotic | β-Lactam Class | $K_{cat}$ ($s^{-1}$) | $K_M$ (µM) | $K_{cat}/K_M$ ($\mu M^{-1} s^{-1}$) | Vmax (µM $min^{-1}$) |
|---|---|---|---|---|---|---|
| SHV-1[25] | Benzylpenicillin | Penicillin | 455 | 20 | 22.8 | - |
| | Cephalothin | Cephalosporin | 10 | 26 | 0.4 | - |
| | Ceftazidime | Cephalosporin | ND | ND | - | - |
| | Cefotaxime | Cephalosporin | ND | ND | - | - |
| | Imipenem | Carbapenem | ND | ND | - | - |
| SHV-2[25] | Benzylpenicillin | Penicillin | NR | 3.8 | NR | 100 |
| | Cephalothin | Cephalosporin | NR | NR | NR | NR |
| | Ceftazidime | Cephalosporin | NR | 24 | NR | 6.5 |
| | Cefotaxime | Cephalosporin | 11 | 18 | 0.6 | 70 |
| | Imipenem | Carbapenem | ND | ND | - | - |
| SHV-38[26] | Benzylpenicillin | Penicillin | 100 | 13 | 7.7 | - |
| | Cephalothin | Cephalosporin | 5 | 100 | 0.05 | - |
| | Ceftazidime | Cephalosporin | 110 | 3800 | 0.03 | - |
| | Cefotaxime | Cephalosporin | 1 | 800 | 0.001 | - |
| | Imipenem | Carbapenem | 0.01 | 24 | 0.0004 | - |

ND - no detected hydrolysis.

NR – not reported.

Data obtained from Liakopoulos *et al.* [25], and Poirel *et al* [26].

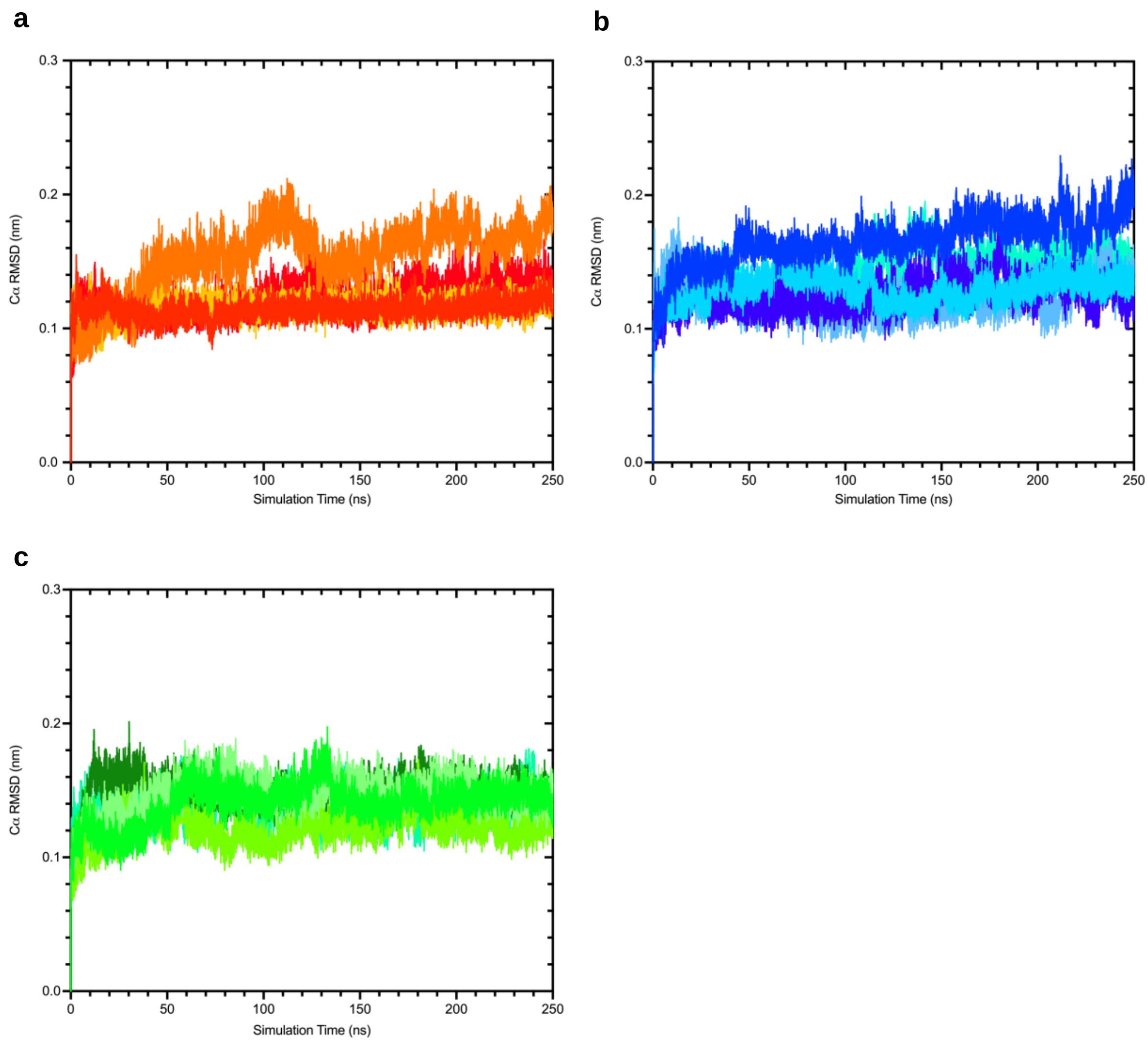


**Figure S3: Temporal evolution of the all-residue average Cα RMSD for 5 repeat equilibrium MD simulations of SHV enzymes.** a) SHV-1, b) SHV-2 and c) SHV-38. The Cα RMSD values were calculated relative to the starting structures. Inspection of frames with larger RMSD values indicated that large fluctuations resulted from the flexibility of the N- and C-termini.

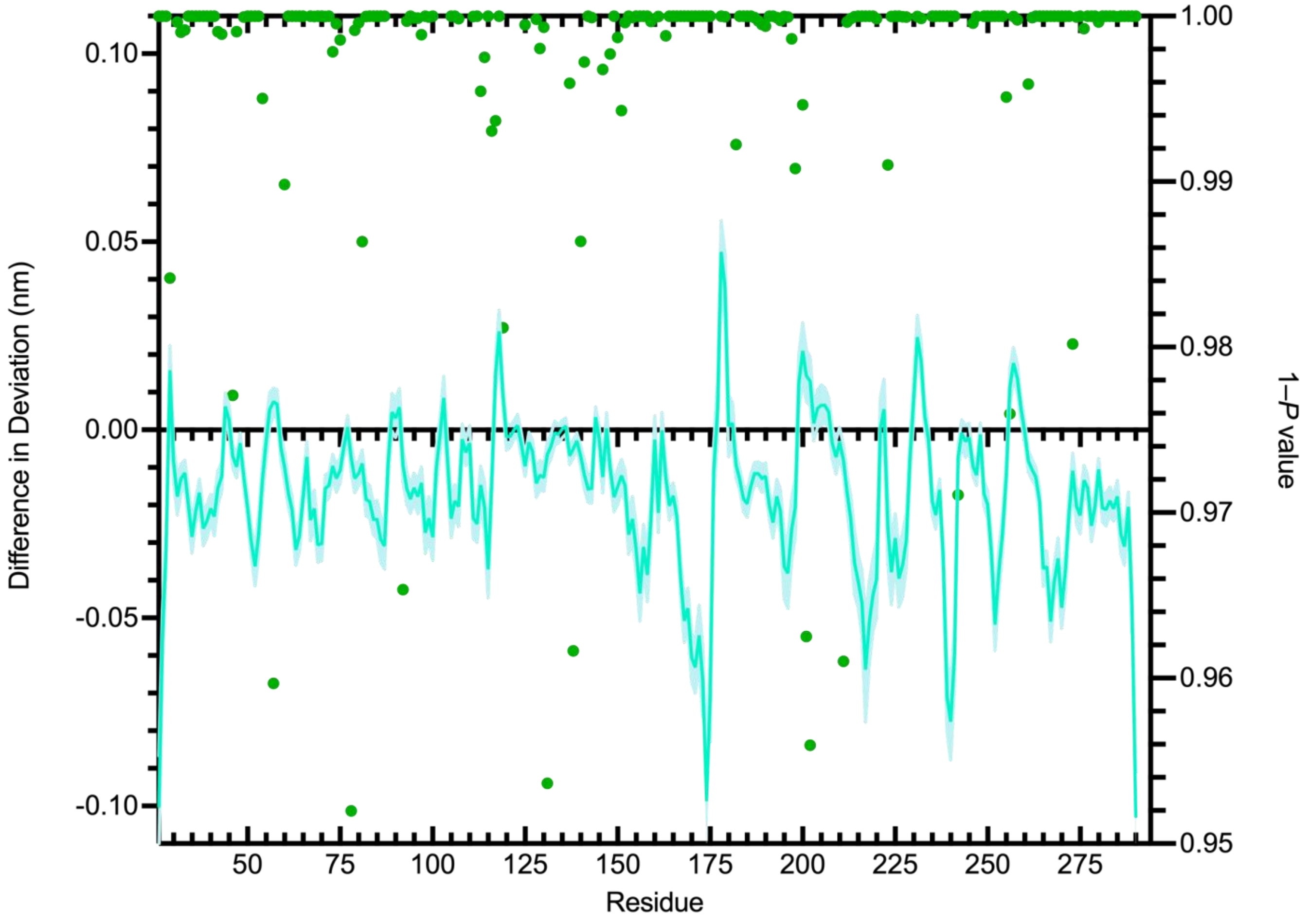


**Figure S4: Differences in residue deviations for SHV-38 vs SHV-2 5ns after perturbation.** Negative values indicate that a residue in SHV-2 deviated more than in SHV-38. Standard errors of the differences are shown in lighter shades above and below the difference line. Green circles denote 1 – *P* values of the difference in deviations between SHV-38 vs SHV-2 Cα deviations (right hand axis).

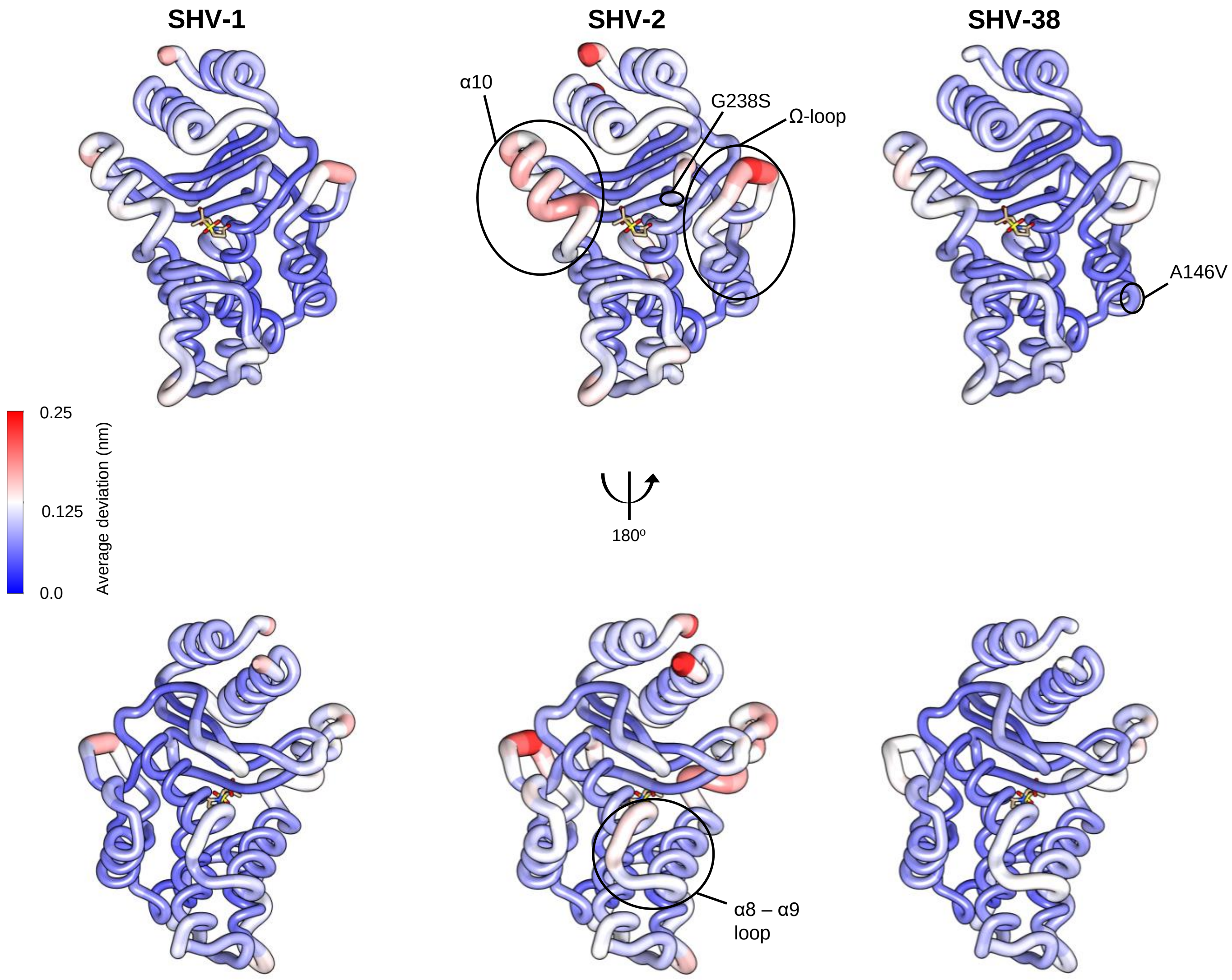


**Figure S5: Average residue deviations of SHV-1, SHV-2 and SHV-38.** Calculated deviations 5 ns after perturbation are rendered onto the SHV-1:sulbactam complex structure (PDB ID 4FH2 [1]). Bound sulbactam is shown as sticks to highlight the active site region. The positions of mutations in SHV-2 (G238S) and SHV-38 (A146V) and regions highlighted for their significant deviations in response to the perturbation in the main text are labelled. Images were generated using Chimera v1.17.3 [27].

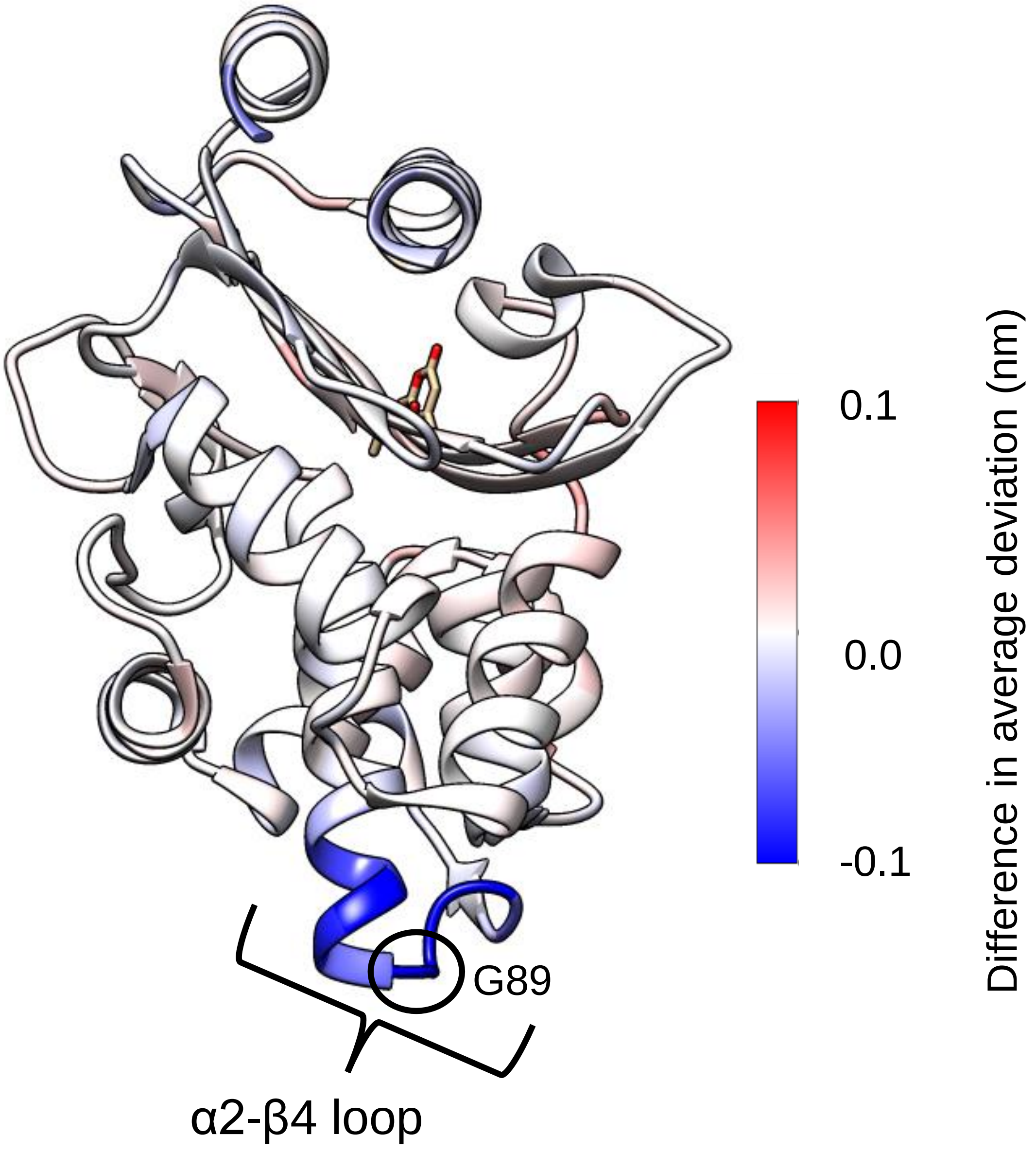


**Figure S6: Average difference in deviation between the null perturbation and the D-NEMD simulations.** A value of 0 indicates no change in the Cα deviation between the null perturbation and the D-NEMD simulations, negative values indicate a reduction in Cα deviation in the D-NEMD simulations compared to the null perturbation and positive values an increase.

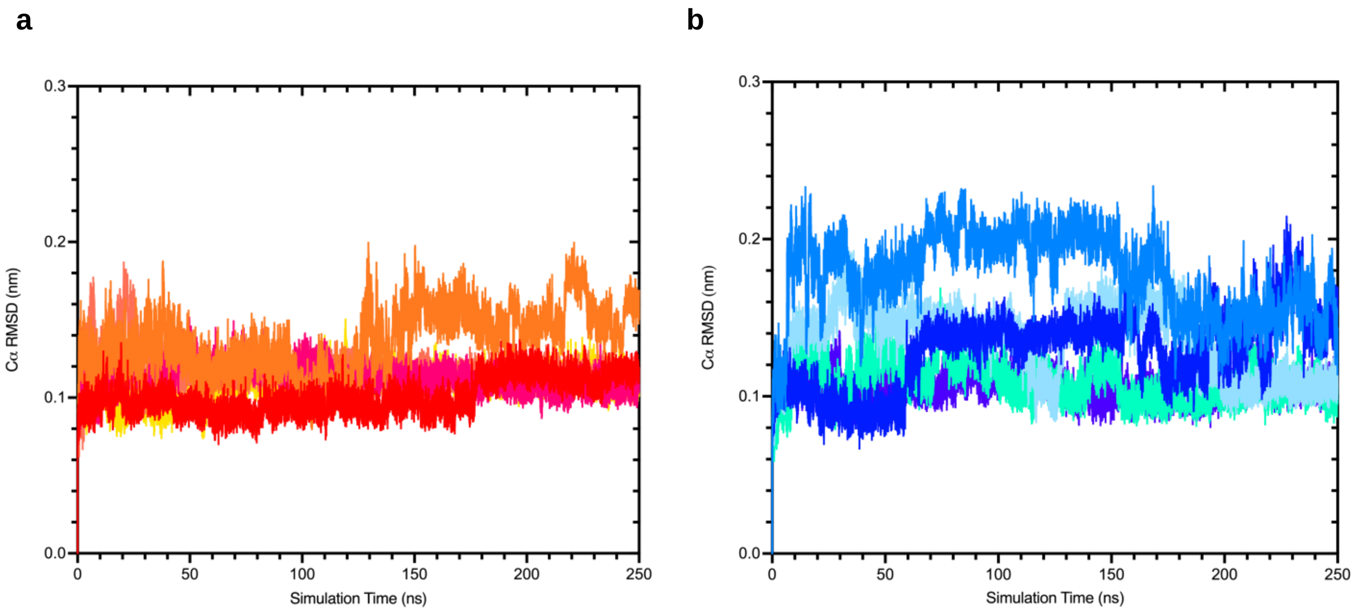


**Figure S7: Temporal evolution of all-residue average Cα RMSD for all 5 repeat equilibrium MD simulations of a) KPC-2 and b) KPC-2$^{G89D}$ enzymes.** Cα RMSD values were calculated relative to the starting structures. Inspection of frames with high RMSD values indicated that large increases in RMSD reflect flexibility of the N- and C-termini.

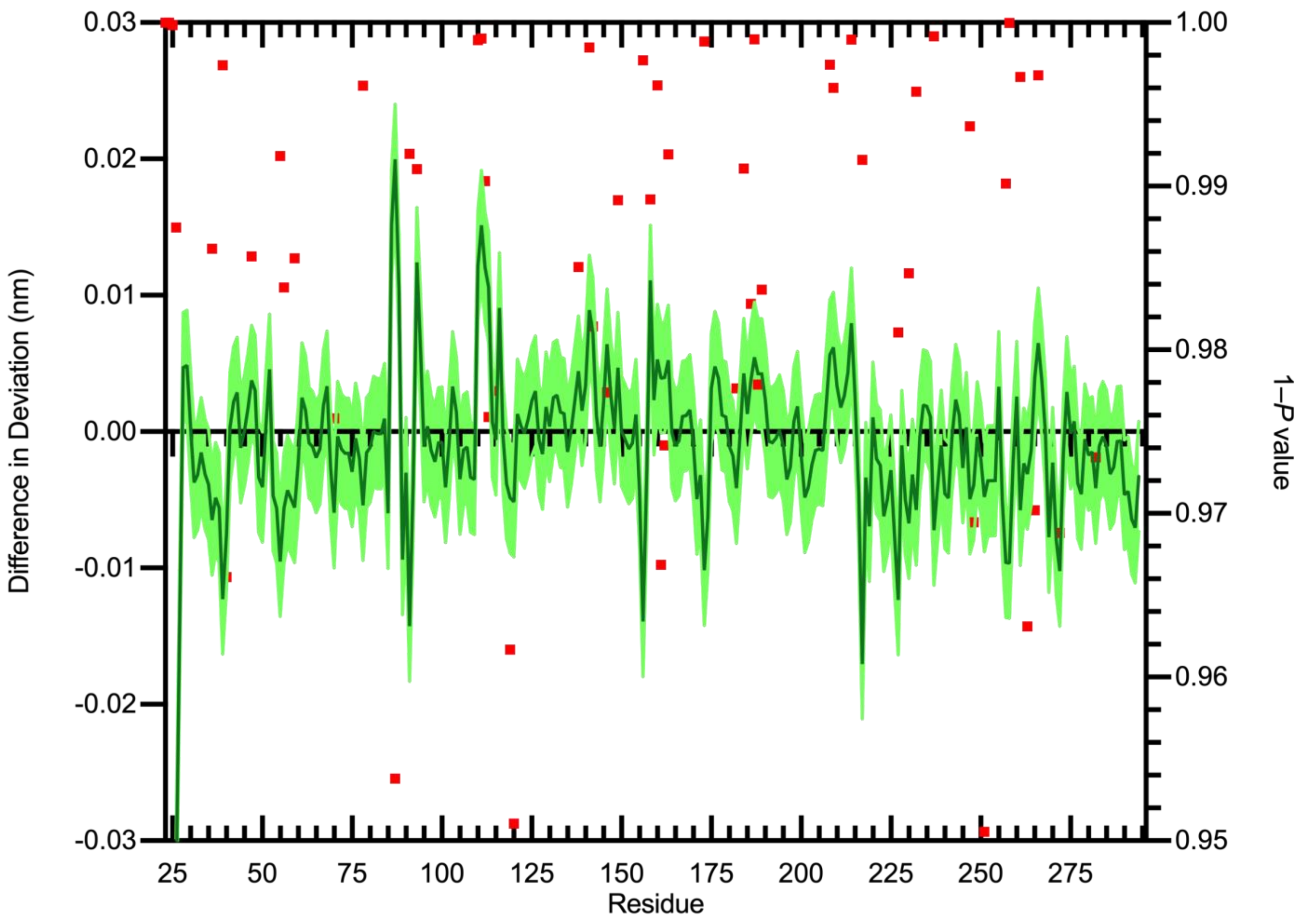


**Figure S8: Differences in average Cα deviation between KPC-2 and KPC-2$^{G89D}$.** Standard error of the difference is shown as pale green shaded region above and below the solid line. Red squares denote 1 – *P* values above 0.95 (indicating statistical significance) for the differences in average Cα deviations between KPC-2 vs. KPC-2$^{G89D}$ (right hand axis).

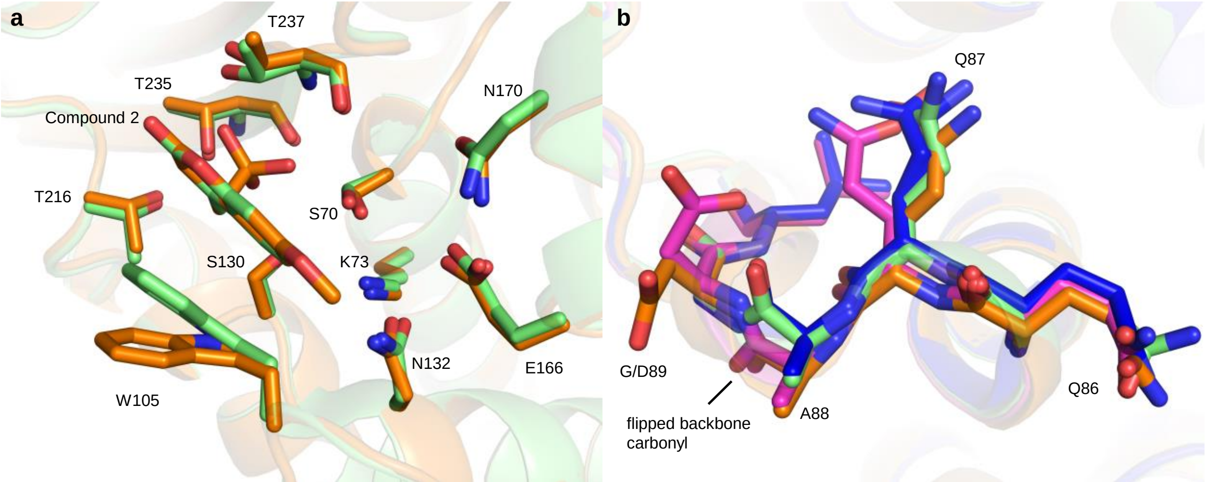


**Figure S9: Validation of the ColabFold[3] KPC-2$^{G89D}$ Model (before minimisation).** a) Active site residues of KPC-2:compound 2 crystal structure (PDB ID 6D16, green) and KPC-2$^{G89D}$ AlphaFold homology model:compound 2 structure (orange). Positions of the heteroaryl phosponate compound 2[5] are also shown. b) The α2-β4 loop (containing residue 89) loop in apo KPC-2$^{G89D}$ (pink), KPC-2$^{G89D}$ AlphaFold homology model:compound 2 complex (orange), KPC-2:compound 2 complex crystal structure (PDB ID 6D16, green) and apo KPC-2 (blue). The AlphaFold homology model determines an accurate estimation of both the active site and α2-β4 loop and correctly captures the flipped backbone carbonyl of alanine 88.

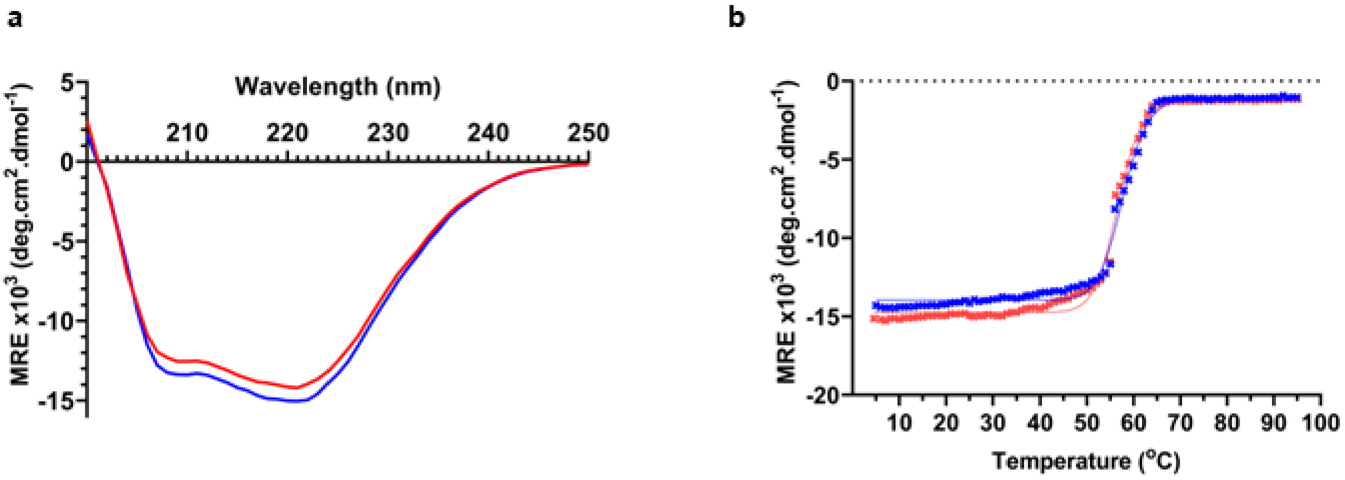


**Figure S10: Circular Dichroism spectroscopy of KPC2 and KPC-2$^{G89D}$.** a) Far-UV circular dichroism spectra of KPC-2 (red) and KPC-2$^{G89D}$ (blue) at a protein concentration of 20 µM and at 25°C. b) Thermal melt experiment (220 nm) of KPC-2 (red, Tm 56.6°C) and KPC-2$^{G89D}$ (blue, Tm 57.7°C). The solid line represents the non-linear regression model used to calculate Tm.

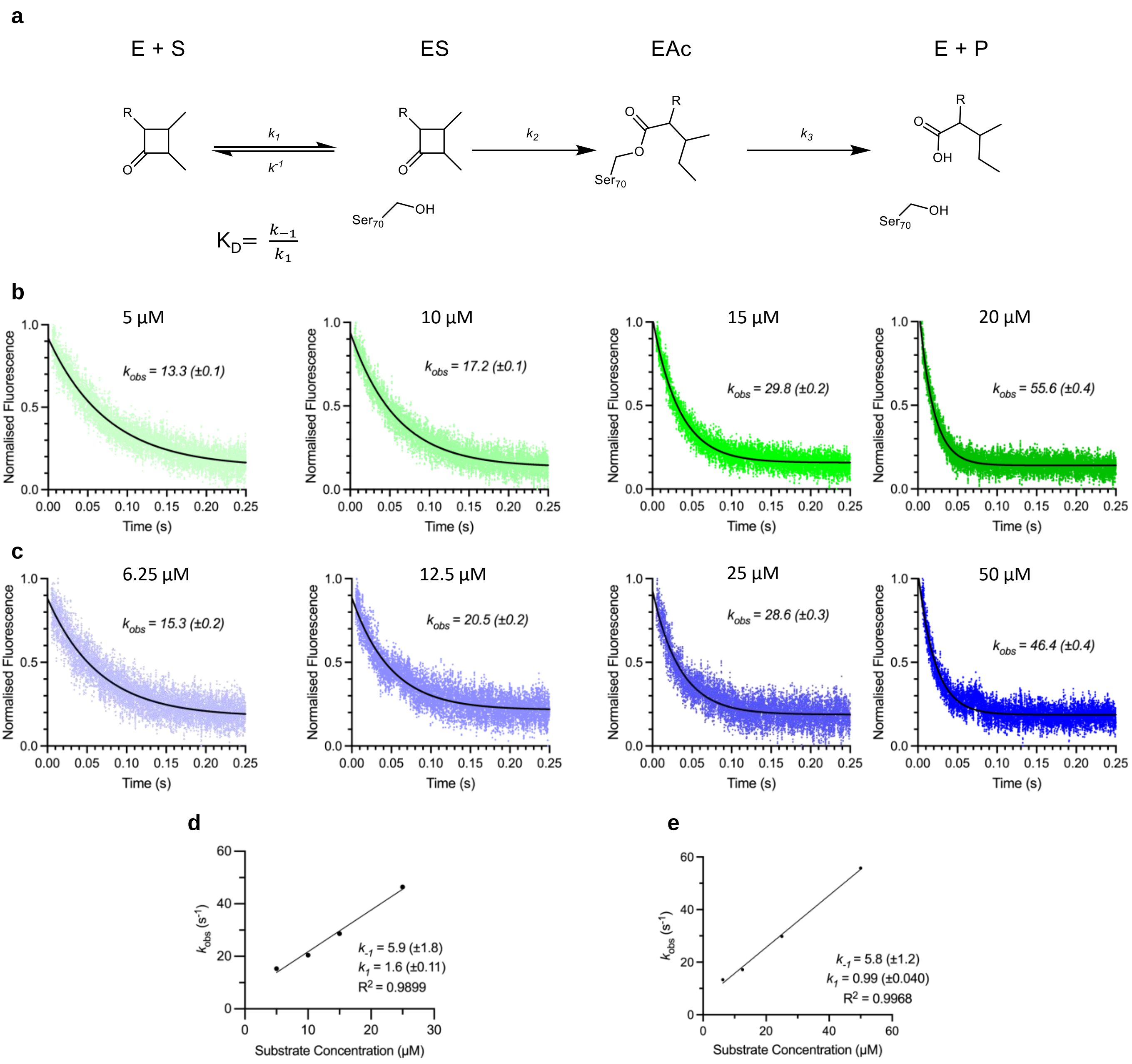


**Figure S11: Binding kinetics of meropenem calculated using pre-steady state kinetics.** a) Kinetic model for binding and hydrolysis of β-lactam substrates by serine β-lactamases. $K_D$ is calculated by $k_{-1}$ / $k_1$. Normalised tryptophan fluorescence curves, measured from 0.005 s to 0.25 s, for meropenem binding to KPC-2 (b) and KPC-2$^{G89D}$ (c) were used to calculate $k_{obs}$ over a range of concentrations using one phase decay curve fitting. $k_{obs}$ was plotted for KPC-2 (d) and KPC-2$^{G89D}$ (e), and a linear regression used to calculate the y-intercept ($k_{-1}$) and the slope ($k_1$). The calculated $K_D$ values are 3.7 µM and 5.9 µM for KPC-2 and KPC-2$^{G89D}$.

**Table S2: Crystallographic data collection and refinement statistics.**

| | KPC-2$^{G89D}$ | KPC-2$^{G89D}$:avibactam | KPC-2$^{G89D/E166Q}$ | KPC-2$^{G89D/E166Q}$:imipenem | KPC-2$^{G89D/E166Q}$:meropenem |
|---|---|---|---|---|---|
| **PDB Code** | 8RWO | 8RWP | 8RWQ | 8RWR | 8RWS |
| **Data Collection** | | | | | |
| Wavelength | 0.81531 | 0.81531 | 0.81531 | 0.81531 | 0.81531 |
| Resolution Range | 59.94-1.13 | 45.39-1.19 | 45.56-1.05 | 45.60-1.03 | 36.39-1.09 |
| Space group | P 21 21 2 | P 21 21 2 | P 21 21 2 | P 21 21 2 | P 21 21 2 |
| Molecules/ASU | 1 | 1 | 1 | 1 | 1 |
| Cell dimensions | | | | | |
| a, b, c (Å) | 59.94 78.62 55.89 | 60.14 78.49 55.64 | 60.36 78.66 55.89 | 60.21 79.45 55.68 | 60.12 78.75 56.16 |
| α, β, γ (°) | 90.00, 90.00, 90.00 | 90.00 90.00 90.00 | 90.00 90.00 90.00 | 90.00 90.00 90.00 | 90.00 90.00 90.00 |
| Multiplicity | 13.2 (12.9) | 13.5 (13.9) | 13.4 (13.3) | 13.6 (13.8) | 13.4 (13.2) |
| Completeness (%) | 100.0 (100.0) | 100.0 (100.0) | 100.0 (99.9) | 98.5 (96.1) | 99.4 (97.4) |
| I/σ(I) | 5.8 (0.5) | 10.0 (0.4) | 8.8 (0.3) | 9.6 (0.3) | 7.0 (0.3) |
| $R_{pim}$ | 0.063 (1.259) | 0.033 (1.005) | 0.036 (1.003) | 0.031 (1.104) | 0.042 (1.105) |
| $CC_{1/2}$ | 0.999 (0.347) | 1.000 (0.295) | 0.999 (0.320) | 1.000 (0.347) | 0.999 (0.324) |
| **Refinement** | | | | | |
| Resolution | 55.89-1.13 | 45.39-1.19 | 45.56-1.05 | 39.73-1.03 | 36.39-1.09 |
| No. reflections | 99264 | 84567 | 124269 | 129928 | 110309 |
| R-work/R-free | 0.1613 / 0.1899 | 0.1554 / 0.1794 | 0.1486 / 0.1688 | 0.1587 / 0.1745 | 0.1740 / 0.1996 |
| No. non-H atoms | | | | | |
| Protein | 2157 | 2142 | 2087 | 2185 | 2200 |
| Solvent | 393 | 382 | 399 | 303 | 306 |
| Ligand | - | 28 | - | 40 | 52 |
| Average B-factors | | | | | |
| Protein | 15.6 | 20.5 | 16.3 | 17.1 | 19.9 |
| Solvent | 32.7 | 38.6 | 36.3 | 35.1 | 34.8 |
| Ligand | - | 20.9 | - | 19.4 | 25.3 |
| R.m.s deviations | | | | | |
| Bond lengths (Å) | 0.007 | 0.008 | 0.006 | 0.007 | 0.007 |
| Bond angles (º) | 0.966 | 0.999 | 0.907 | 1.175 | 1.019 |
| Ramachandran (%) | | | | | |
| Outliers | 0.00 | 0.00 | 0.00 | 0.00 | 0.00 |
| Favoured | 98.88 | 98.88 | 98.88 | 98.88 | 98.49 |

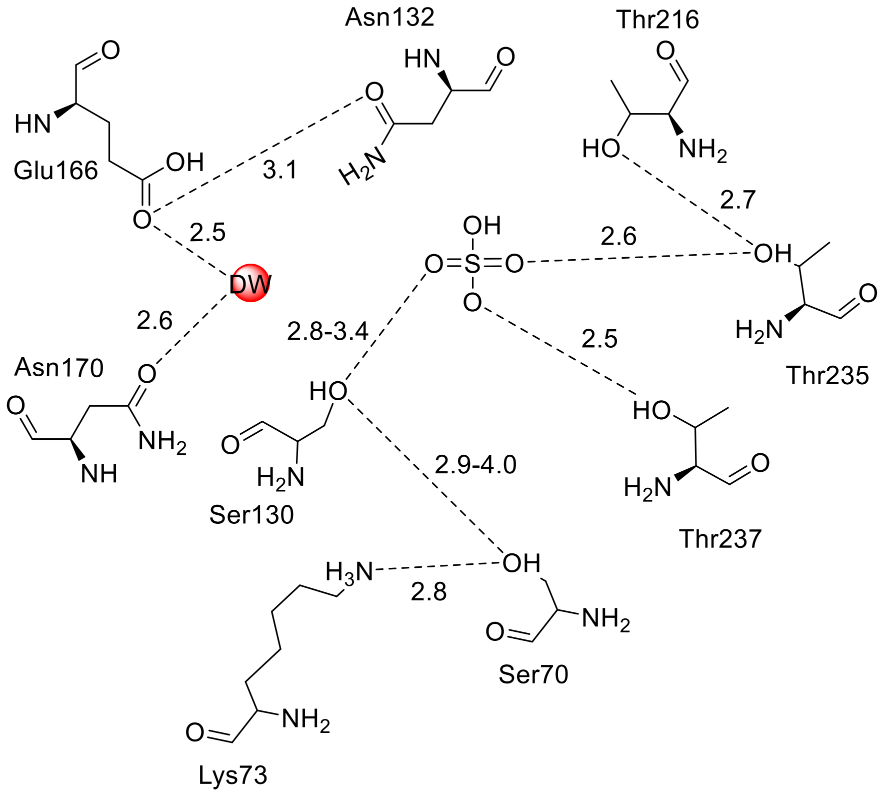


**Figure S12: Active Site Interaction Diagram of uncomplexed KPC-2$^{G89D}$.** Hydrogen bonding interactions are highlighted (black dashed line) with interatomic distances shown in Å. Distance ranges are given for hydrogen bonding around Ser130 as the Ser130 side chain was modelled in two conformations.

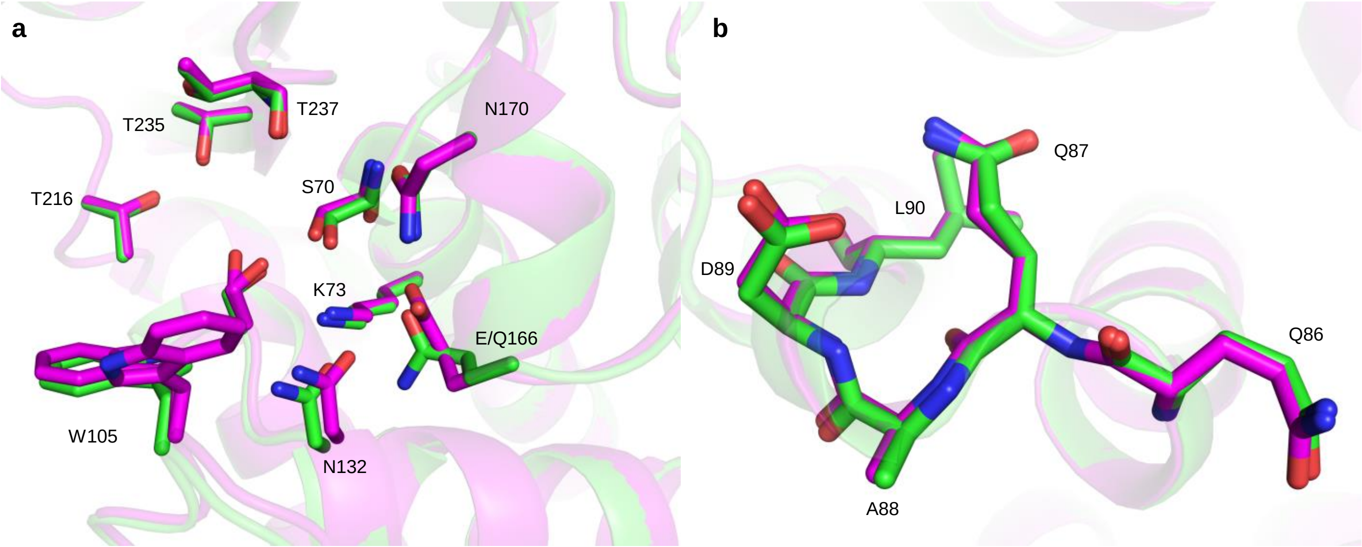


**Figure S13: Structure of Uncomplexed KPC-2$^{G89D/E166Q}$.** a) Active site residues of KPC-2$^{G89D}$ (pink) and KPC-2$^{G89D/E166Q}$. b) Alignment of the α2-β4 loop in KPC-2$^{G89D}$ and KPC-2$^{G89D/E166Q}$ structures.

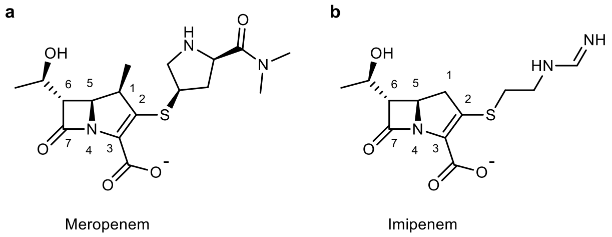


**Figure S14: Atom numbering of meropenem and imipenem**. The β-lactam carbonyl carbon (C-7) is the position of nucleophilic attack by Ser70, breaking the scissile β-lactam amide bond and forming the acyl-enzyme complex. Meropenem (a) and imipenem (b) differ in their C2 substituents and the presence of the meropenem 1β-methyl group (cf. imipenem 1β-hydrogen).

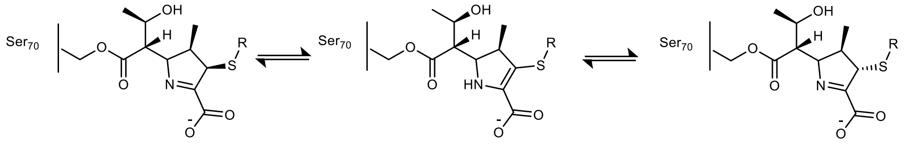


**Figure S15. Carbapenem acylenzyme tautomerisation.** In the carbapenem acyl-enzyme complex, the double bond in the pyrroline ring can migrate from C3 = C2 (Δ2, enamine) to N4 = C3 (Δ1, imine, in 2*R*- and 2*S*- configurations), resulting in three possible forms, Δ2, Δ1-(2*R*) and Δ1-(2*S*).

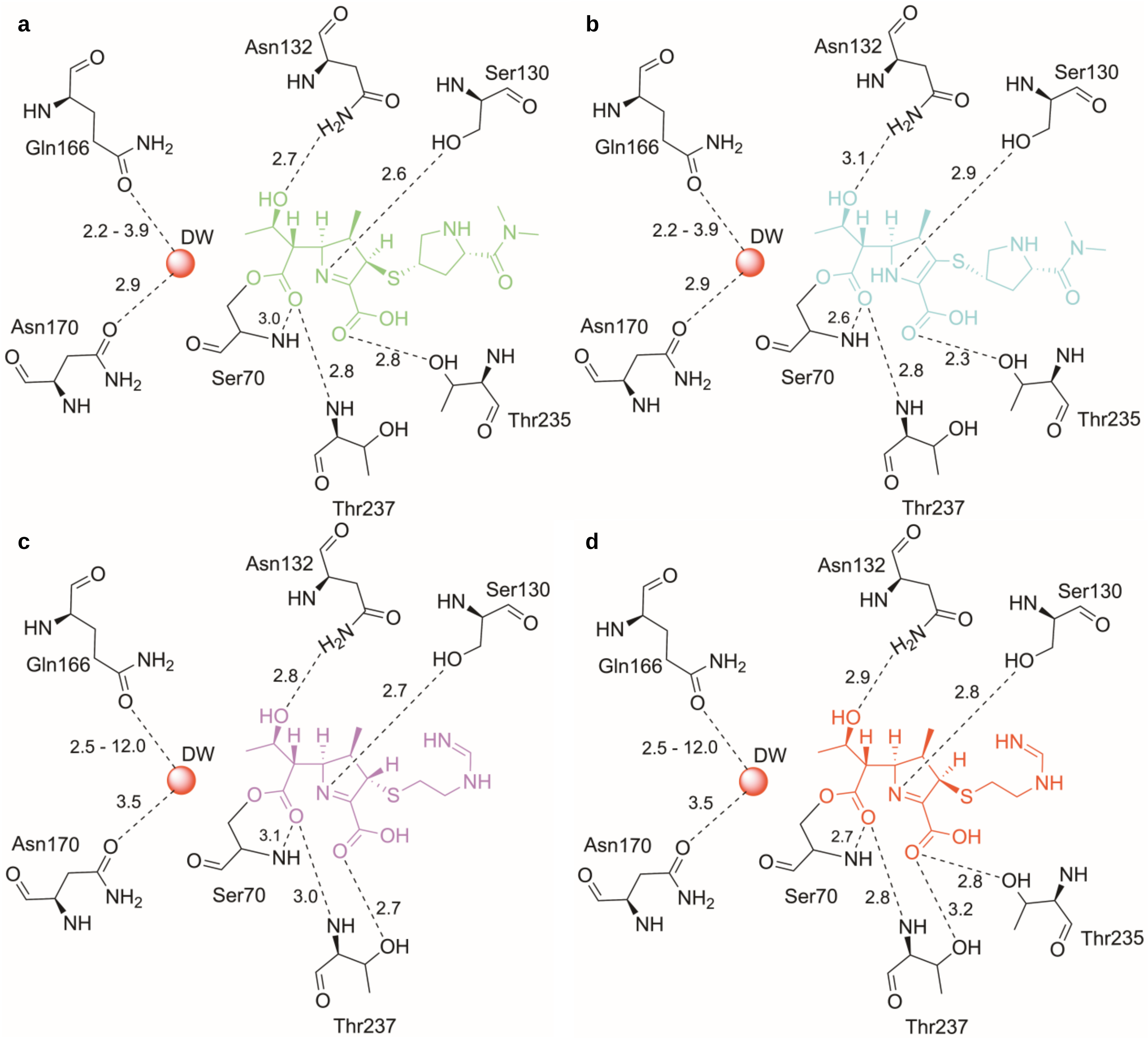


**Figure S16. Ligand interactions of KPC-2$^{G89D}$:carbapenem complexes.** a) KPC-2$^{G89D/E166Q}$:meropenem Δ1-(2*R*) tautomer. b) KPC-2$^{G89D/E166Q}$:meropenem Δ2 tautomer. c) KPC-2$^{G89D/E166Q}$:imipenem Δ1-(2*S*) tautomer. d) KPC-2$^{G89D/E166Q}$:imipenem Δ1-(2*R*) tautomer. Ranges of values for the Gln166-OE1 - deacylating water (DW) indicate minimum and maximum distances observed for multiple conformations modelled in the crystal structures. The larger distance range for imipenem reflects the modelling of Gln166 in the 'out' conformation in some conformers of the imipenem-derived structure; i.e. with the side chain pointing into bulk solvent rather than towards DW. Only 'in' conformers of Gln166 were modelled in the meropenem-derived structures. Structures were generated using LigPlot+ v2.2 [28].

**a** **b**

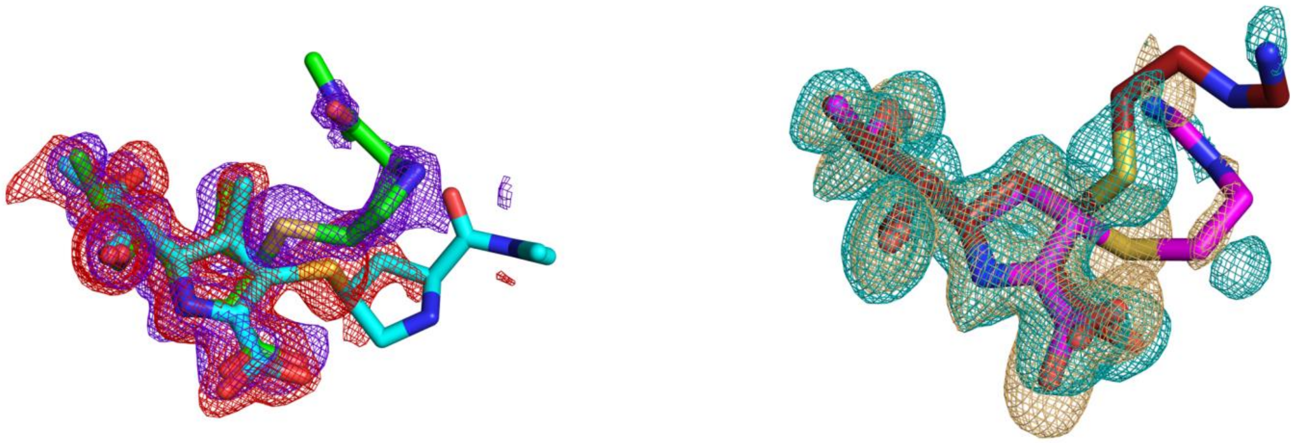

**Figure S17: Omit maps of KPC-2G89D/E166Q:carbapenem complexes.** Modelled ligands with *F*o-*F*c omit density calculated after ligand removal contoured at 3σ. a) Δ1-(2*R*) (ligand carbon atoms green, blue density) and Δ2 (ligand carbons cyan, red density) tautomers of the meropenem-derived acylenzyme. b) Δ1-(2*R*) (red ligand carbons red, cyan density) and Δ1-(2*S*) (magenta ligand, orange density) tautomers of the imipenem-derived acylenzyme. C2 substituents are known to be flexible and make few/no direct electrostatic contacts with enzyme residues, reflected in the poor electron density resolution after the sulphur atom.

**a**

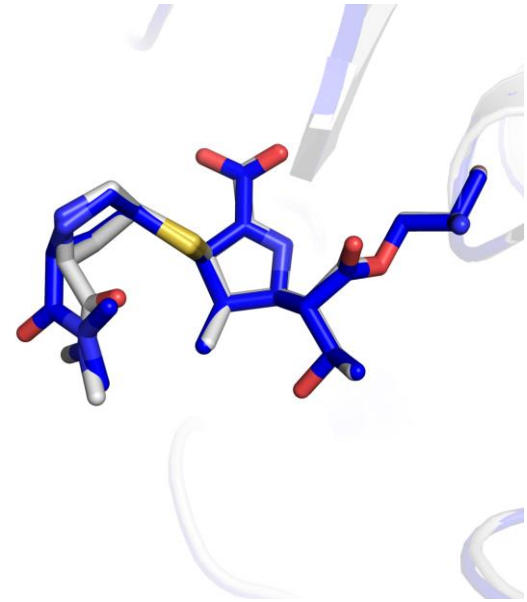

**b**

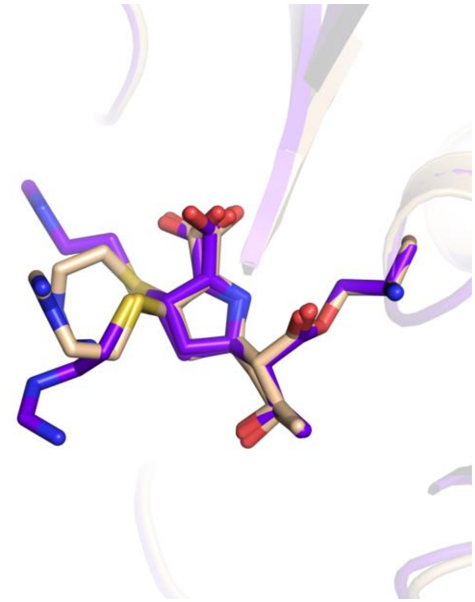

**Figure S18: Carbapenem orientation in the KPC-2$^{G89D/E166Q}$ active sites.** Structural alignment between KPC-2$^{G89D/E166Q}$:meropenem and KPC-2$^{E166Q}$:meropenem (PDB 8AKL) (a) and between KPC-2$^{G89D/E166Q}$:imipenem and KPC-2$^{E166Q}$:imipenem (PDB 8AKK). The C2 substituent (Figure S14) is known to make few/no interactions with enzyme residues and subsequently is highly flexible, particularly after the sulphur atom. This accounts for the difference in binding conformation in this region of the carbapenem, particularly for the imipenem molecule (b).

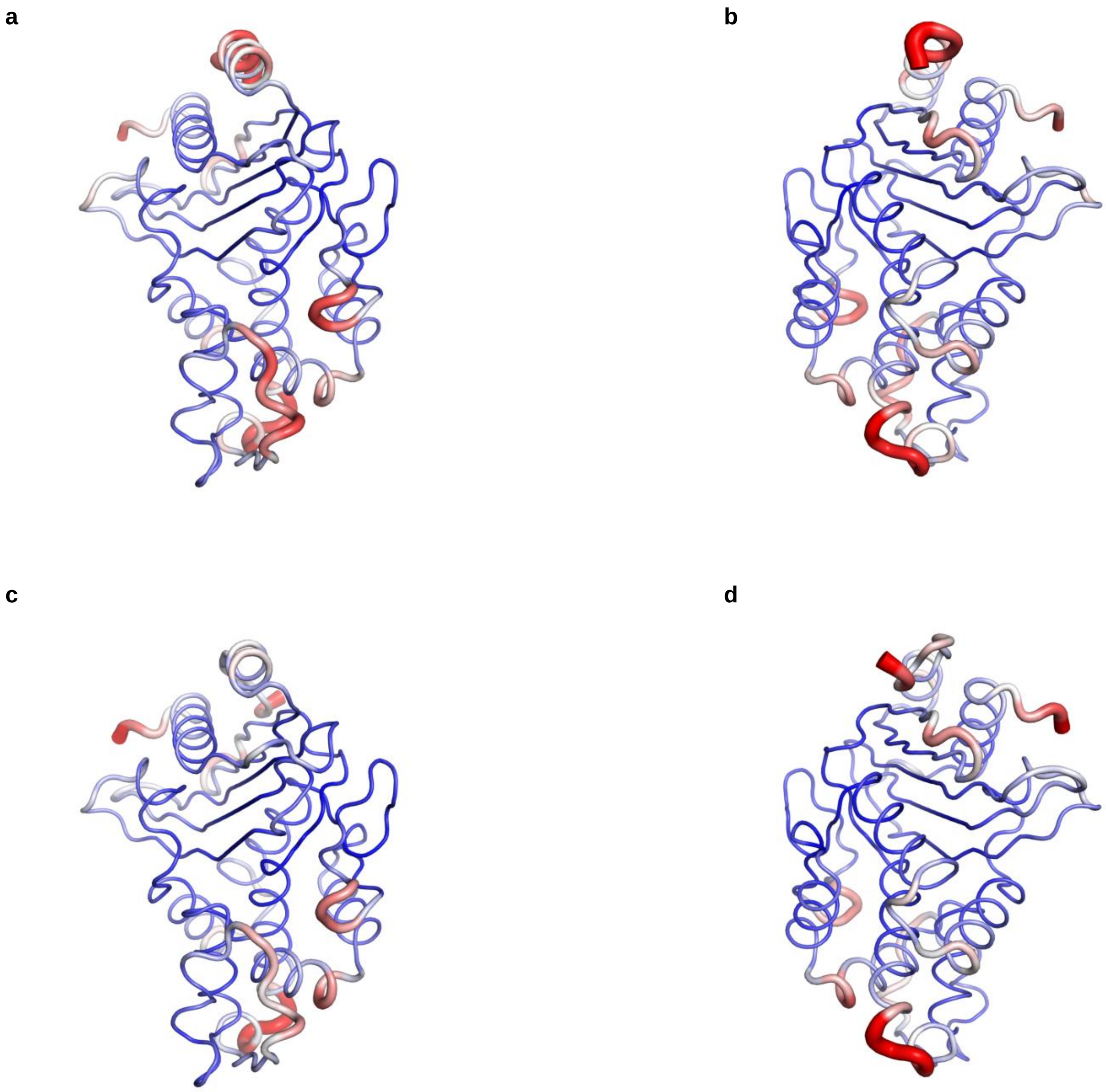


**Figure S19: B-factor analysis of the KPC-2$^{G89D/E166Q}$:carbapenem complexes.** B-factors rendered onto KPC-2$^{G89D/E166Q}$:meropenem (a-b) and KPC-2$^{G89D/E166Q}$:imipenem (c-d) shows high B-factors for both the Ω- and α4-β2 loops. The scale represents B-factor values between 10 (thin, blue) to 35 (thick, red).

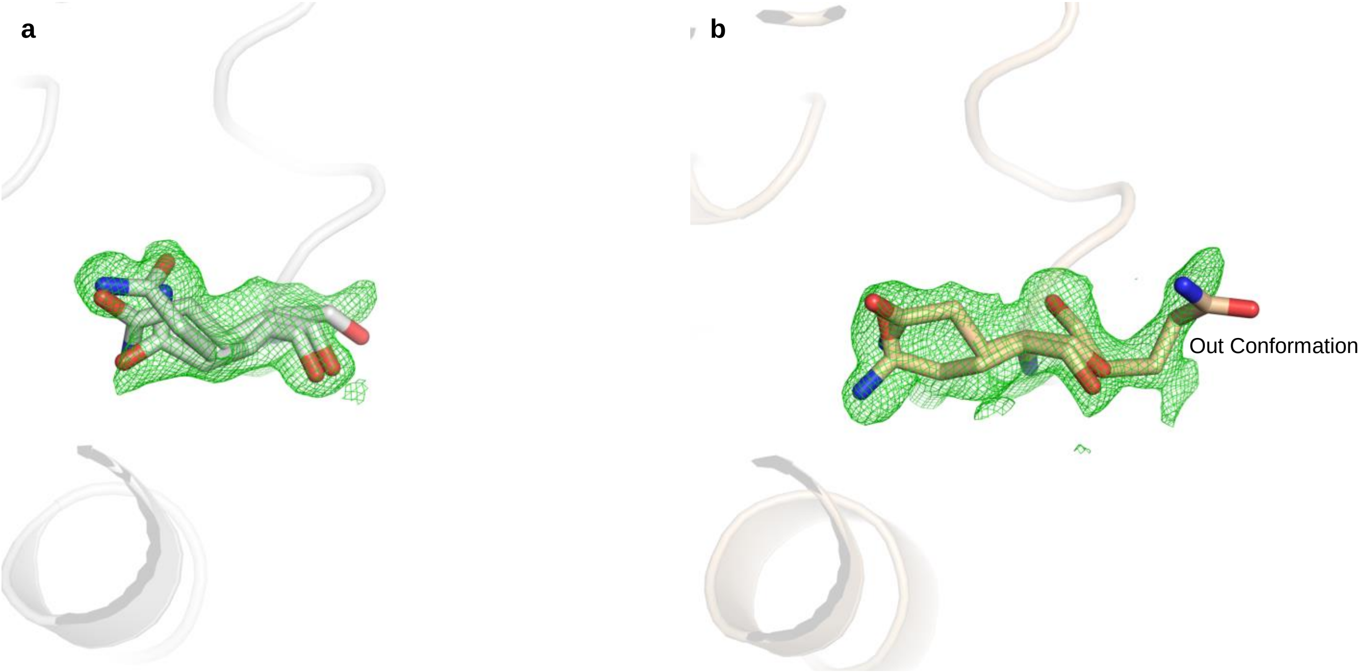


**Figure S20: Polder map of Gln166 in KPC-2$^{G89D/E166Q}$:carbapenem complexes.** $F_o$-$F_c$ polder maps contoured at 3σ. a) Gln166 in the KPC-2$^{G89D/E166Q}$:meropenem complex. b) Gln166 in the KPC-2$^{G89D/E166Q}$:imipenem complex, showing the 'out' conformation.

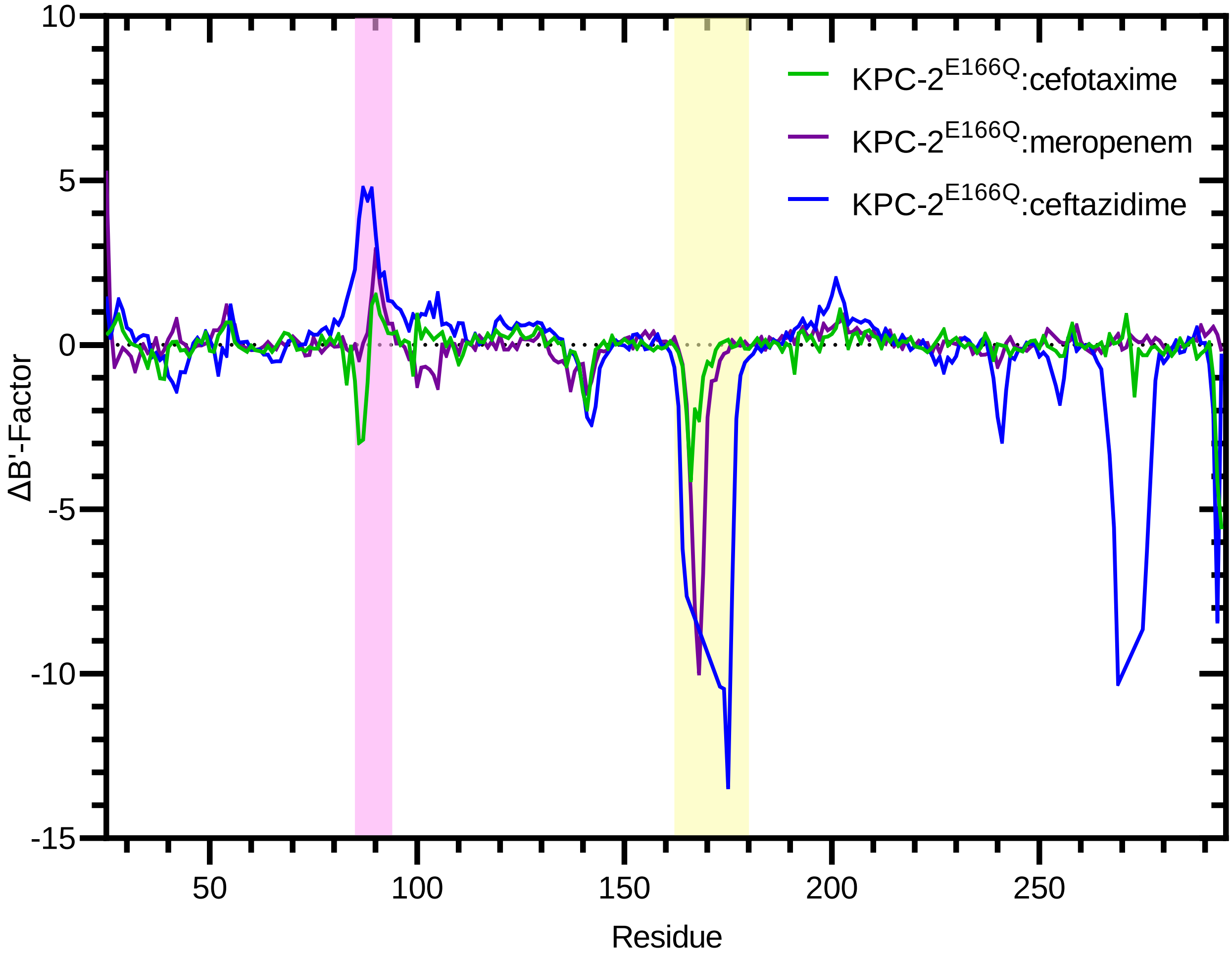


**Figure S21: Difference in adjusted B-factors (B'-factors) between uncomplexed KPC-2 and KPC-2$^{E166Q}$:β-lactam acyl-enzyme complexes.** Plot shows differences in B'-factors between uncomplexed KPC-2 (PDB 5UL8) and: KPC-2$^{E166Q}$:cefotaxime (PDB ID 6Z23[16], green); KPC-2 $^{E166Q}$:meropenem (PDB 8AK1[24], purple) and KPC-2$^{E166Q}$:ceftazidime (PDB ID 6Z24[16], blue). Negative values indicate residues for which the B'-factor of the respective KPC-2$^{E166Q}$:β-lactam complex is greater than that of uncomplexed KPC-2. Turnover is slowest for ceftazidime ($k_{cat}$ = 1.9 $s^{-1}$) and fastest for cefotaxime ($k_{cat}$ = 76 $s^{-1}$), demonstrating that low B'-factors for the α2 - β4 loop (pink highlight) and high B'-factors for Ω-loop (yellow highlight) in the respective acylenzymes correlate well with turnover of specific substrates by KPC-2.

**Table S3: DW occupancy and B-factor in KPC-2$^{G89D/E166Q}$:carbapenem complexes.**

| Structure | DW Occupancy | DW B-factor |
|---|---|---|
| KPC-2$^{G89D/E166Q}$:meropenem | 0.58 | 24.49 |
| KPC-2$^{G89D/E166Q}$:imipenem | 1.00 | 19.74 |

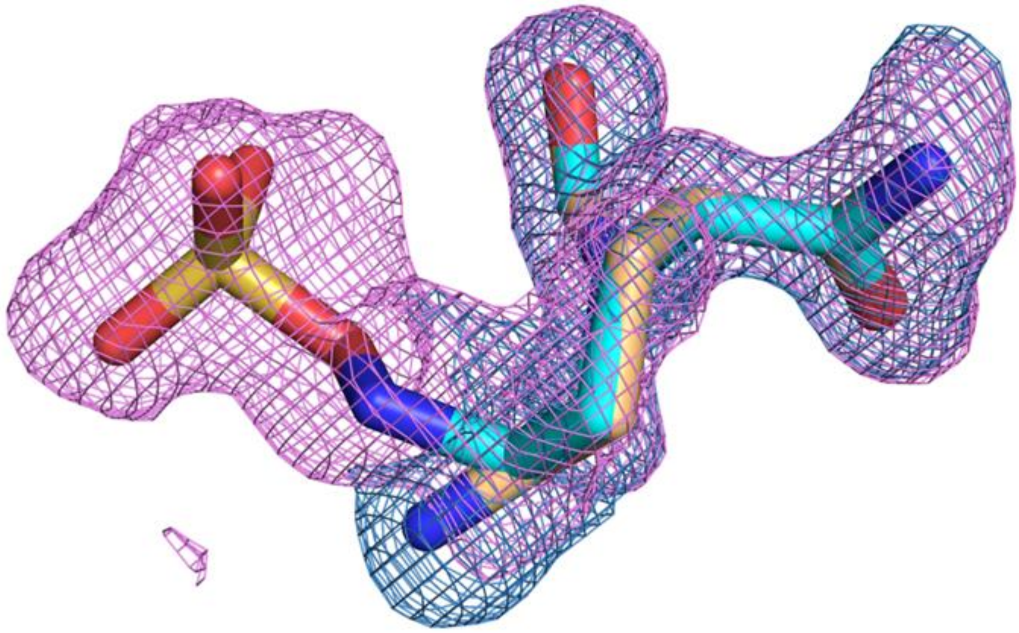

**Figure S22**: **Omit map density of KPC-2$^{G89D}$:avibactam complex.** Omit map density contoured at 3σ for sulphated (magenta mesh) and desulphated (teal mesh) avibactam structures modelled.

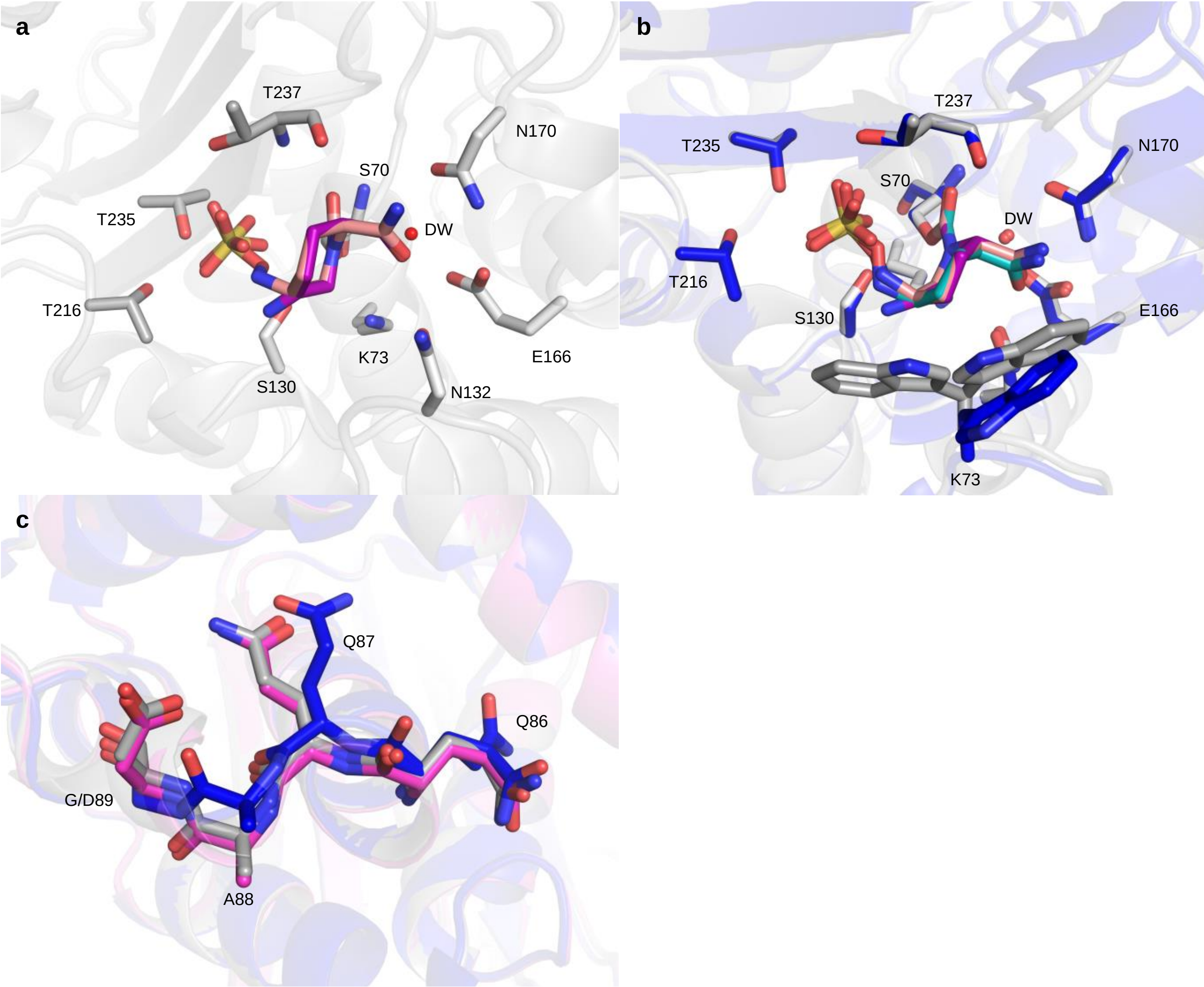


**Figure S23: KPC-2$^{G89D}$:avibactam structure.** a) Active site of KPC-2$^{G89D}$ in complex with avibactam. Both the sulphated (orange) and desulphated (purple) forms of avibactam could be modelled. Both the sulphated and desulphated forms were refined to an occupancy of 0.5 each. b) KPC-2$^{G89D}$:avibactam complex structure (KPC-2$^{G89D}$ side chain carbon atoms grey, ligands purple and orange) aligned with KPC-2:avibactam (carbon atoms blue, ligand cyan, PDB ID 4ZBE [29]). Overall RMSD between the two complexes is 0.2 Å. c) Alignment of KPC-2$^{G89D}$:avibactam (grey), KPC-2:avibactam (blue) and uncomplexed KPC-2$^{G89D}$ (pink), showing the α2-β4 loop.

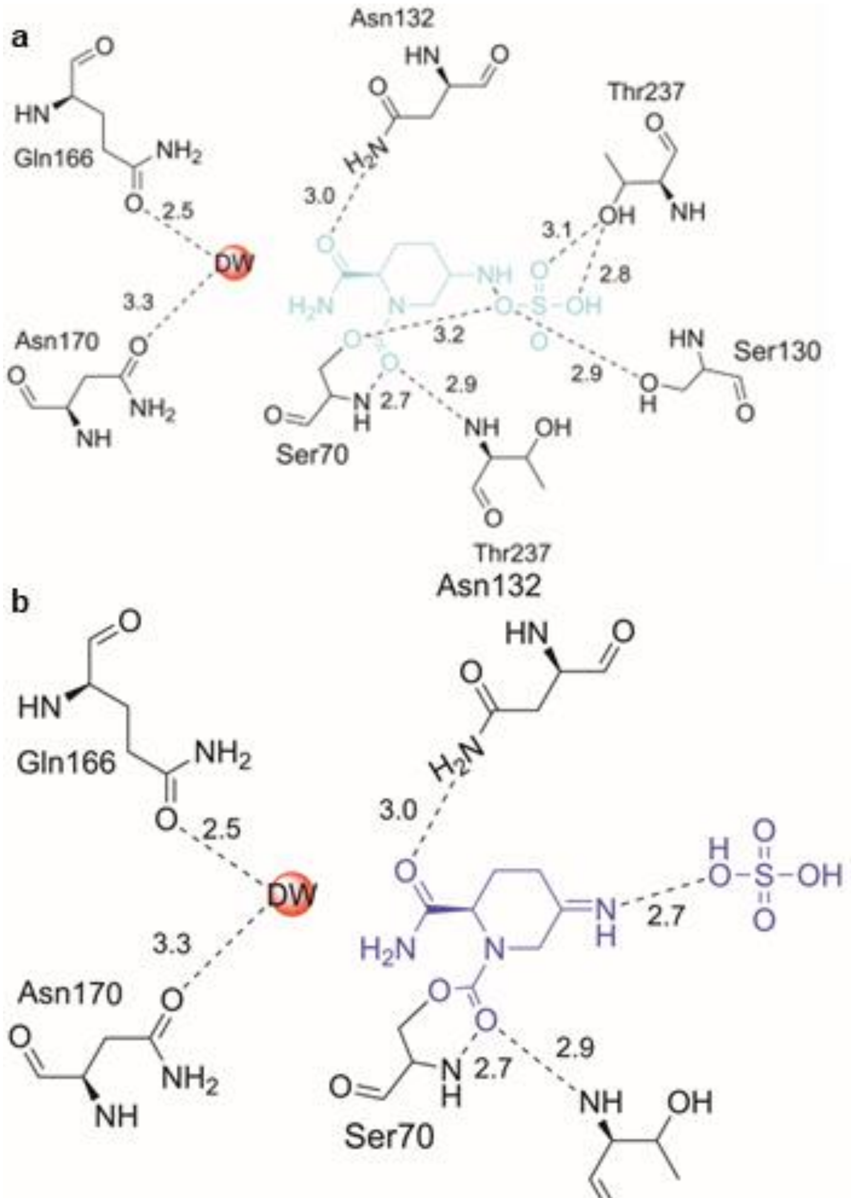


**Figure S24: Ligand interactions of the KPC-2$^{G89D}$:avibactam complex**. Hydrogen bonds are shown as dashed lines, distances in Å. The avibactam carbamyl is coloured cyan (with sulphate present, a) or purple (desulphated, b).

SUPPLEMENTARY INFORMATION REFERENCES


1. Rodkey, E.A., et al., *Crystal Structure of a Preacylation Complex of the β-Lactamase Inhibitor Sulbactam Bound to a Sulfenamide Bond-Containing Thiol-β-lactamase.* Journal of the American Chemical Society, 2012. **134**(40): p. 16798-16804.
2. Kuzin, A.P., et al., *Structure of the SHV-1 β-Lactamase.* Biochemistry, 1999. **38**(18): p. 5720-5727.
3. Mirdita, M., et al., *ColabFold: making protein folding accessible to all.* Nature Methods, 2022. **19**(6): p. 679-682.
4. Olsson, M.H.M., et al., *PROPKA3: Consistent Treatment of Internal and Surface Residues in Empirical pKa Predictions.* Journal of Chemical Theory and Computation, 2011. **7**(2): p. 525-537.
5. Pemberton, O.A., et al., *Heteroaryl Phosphonates as Noncovalent Inhibitors of Both Serine- and Metallocarbapenemases.* Journal of Medicinal Chemistry, 2019. **62**(18): p. 8480-8496.
6. Emsley, P., et al., *Features and development of Coot.* Acta Crystallogr D Biol Crystallogr, 2010. **66**(Pt 4): p. 486-501.
7. Bauer, P., B. Hess, and E. Lindahl, *GROMACS 2019.1 Source Code*. 2019, Zenodo.
8. Tian, C., et al., *ff19SB: Amino-Acid-Specific Protein Backbone Parameters Trained against Quantum Mechanics Energy Surfaces in Solution.* Journal of Chemical Theory and Computation, 2020. **16**(1): p. 528-552.
9. Wang, J., et al., *Development and testing of a general amber force field.* J Comput Chem, 2004. **25**(9): p. 1157-74.
10. Sousa da Silva, A.W. and W.F. Vranken, *ACPYPE - AnteChamber PYthon Parser interfacE.* BMC Research Notes, 2012. **5**(1): p. 367.
11. Vanquelef, E., et al., *R.E.D. Server: a web service for deriving RESP and ESP charges and building force field libraries for new molecules and molecular fragments.* Nucleic Acids Research, 2011. **39**(suppl_2): p. W511-W517.
12. Jorgensen, W., J. Chandrasekhar, and J. Madura, *Impey, RW; Klein, ML.* J. Chem. Phys, 1983. **79**: p. 926.
13. Berendsen, H.J.C., et al., *Molecular dynamics with coupling to an external bath.* The Journal of Chemical Physics, 1984. **81**(8): p. 3684-3690.
14. Parrinello, M. and A. Rahman, *Polymorphic transitions in single crystals: A new molecular dynamics method.* Journal of Applied Physics, 1981. **52**(12): p. 7182-7190.
15. Oliveira, A.S.F., et al., *Dynamical nonequilibrium molecular dynamics reveals the structural basis for allostery and signal propagation in biomolecular systems.* The European Physical Journal B, 2021. **94**(7): p. 144.
16. Tooke, C.L., et al., *Natural variants modify Klebsiella pneumoniae carbapenemase (KPC) acyl–enzyme conformational dynamics to extend antibiotic resistance.* Journal of Biological Chemistry, 2021. **296**: p. 100126.
17. Tooke, C.L., et al., *Molecular Basis of Class A β-Lactamase Inhibition by Relebactam.* Antimicrob Agents Chemother, 2019. **63**(10).
18. Cantu, C., III and T. Palzkill, *The Role of Residue 238 of TEM-1 β-Lactamase in the Hydrolysis of Extended-spectrum Antibiotics *.* Journal of Biological Chemistry, 1998. **273**(41): p. 26603-26609.
19. Queenan, A.M., et al., *Hydrolysis and inhibition profiles of beta-lactamases from molecular classes A to D with doripenem, imipenem, and meropenem.* Antimicrobial agents and chemotherapy, 2010. **54**(1): p. 565-569.
20. Hutchins, G.H., et al., *Precision design of single and multi-heme de novo proteins.* bioRxiv, 2020: p. 2020.09.24.311514.
21. Winter, G., et al., *DIALS: implementation and evaluation of a new integration package.* Acta crystallographica. Section D, Structural biology, 2018. **74**(Pt 2): p. 85-97.
